# A generative deep learning method for global species distribution prediction

**DOI:** 10.1101/2024.12.10.627845

**Authors:** Yujing Yan, Bin Shao, Jiawei Yan, Charles C. Davis

**Affiliations:** Department of Organismic and Evolutionary Biology, Harvard University Herbaria, Harvard University, Cambridge MA 02138, USA; Department of Molecular and Cellular Biology, Harvard University, Cambridge MA 02138, USA; Independent researcher, 100 N Gushan Rd, Shanghai, 200135, China

## Abstract

Anthropogenic pressures on biodiversity necessitate efficient and scalable methods to predict global species distributions. Current species distribution models (SDMs) face limitations with large-scale datasets, complex interspecies interactions, and data quality. Here, we introduce EcoVAE, an autoencoder-based generative model that integrates bioclimatic variables with georeferenced occurrences. The model is trained separately for plants, butterflies, and mammals to predict global distributions at both genus and species levels. EcoVAE achieves high precision and speed, outperforming traditional SDMs in spatial block cross-validation. Through unsupervised learning, it captures underlying distribution patterns and reveals species associations that align with known prey-predator relationships. Additionally, it evaluates global sampling efforts and interpolates distributions in data-limited regions, offering new applications for biodiversity exploration and monitoring.

## Introduction

Anthropogenic pressures have intensified the need for efficient and scalable methods to predict species distributions for attaining a clearer understanding of biodiversity. Over the past three decades, species distribution modeling (SDM) has become an essential tool for this purpose^1–5^, typically using species occurrence data and environmental variables to predict distributions through statistical and machine learning (ML) algorithms^6–9^. While the mobilization of vast amounts of specimen records and the rapid accumulation of observational data have greatly promoted the development of SDMs^10–12^, several challenges remain.

First, current SDMs struggle to handle large-scale datasets in our big data era^10–14^, especially for modeling species assemblages. Traditional methods usually address these tasks by fitting single-species SDMs to environmental predictors separately and then combined them to derive community outputs^15–17^. Second, most SDMs were primarily developed based on the climate equilibrium assumption and overlook complex interspecies interactions, limiting their ecological relevance and utility in modeling community dynamics^16,18–20^. Recently developed methods, such as joint species distribution models (JSDMs) and explicit multi-species models, address this gap by simultaneously modeling species abundances or occurrences, capturing residual correlations between taxa and responses to environmental variables^21–23^. However, these models remain computationally intensive and exhibit limited scalability. This critical issue prevents them from leveraging the massive and valuable datasets available via platforms like GBIF and eBird^24^. Finally, reliance on environmental variables introduces additional issues for traditional SDMs, including collinearity and limited availability in certain regions^25,26^, further constraining model accuracy and applicability.

Generative deep learning models are designed to learn and reproduce complex, nonlinear relationships within data, enabling the generation of new, realistic data instances. They have been widely adopted in various fields, including natural language processing^27^, image generation^28^, data capturing^29^, and medicine^30^. Among these models, autoencoders are designed to compress and reconstruct data in an unsupervised way, making it effective for data denoising, interpolation, and handling randomly missing data^31^. The Variational Autoencoder (VAE) employs a probabilistic latent space, and has been applied to study spatial correlation structures among bird species^32^ and to support individual based ecosystem modeling^33^.

Here, we present for the first time an autoencoder-based framework, named Ecological Variational Autoencoder (EcoVAE), to predict species distributions using large-scale presence-only and climatic data (Fig. 1a). We present two versions of EcoVAE: one learns the patterns of global species distributions using occurrences only (EcoVAE-o); the other uses occurrences and climatic variables together (EcoVAE-c).

**Figure 1.**
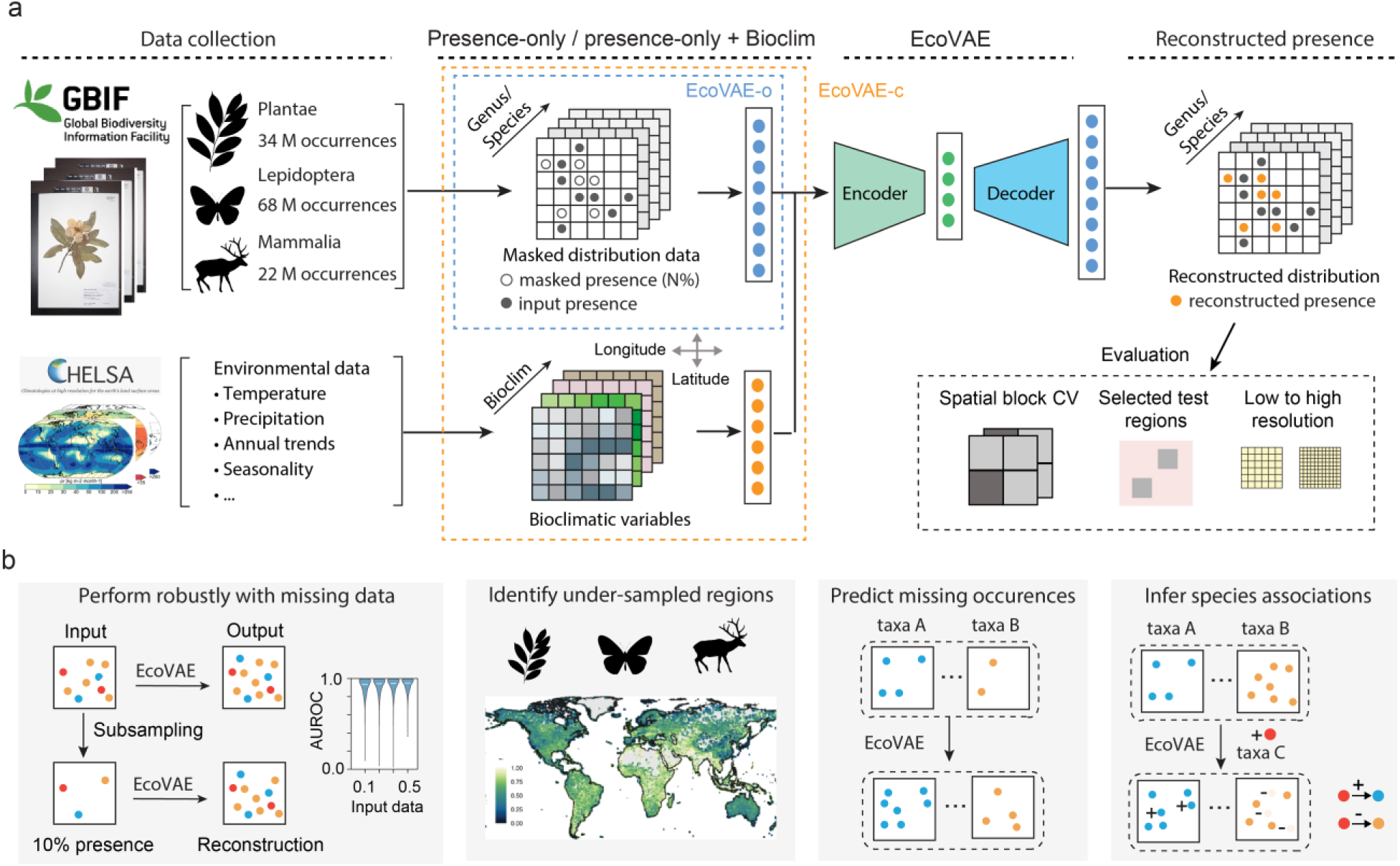
Schematic representation of the model training and application pipeline. **a,** Training of the EcoVAE model. We developed two versions of the EcoVAE model: EcoVAE-o (presence-only inputs) and EcoVAE-c (presence plus bioclimatic covariates as input). **b,** Applications of the EcoVAE model.

To demonstrate the effectiveness of our framework, we trained customized EcoVAE models on a massive global dataset including nearly 34 million georeferenced vouchered occurrences (positive detection records) from the GBIF platform spanning 11,555 plant genera and 127,281 species. Both EcoVAE-o and EcoVAE-c efficiently process this large-scale dataset with fast computational times and accommodates varying rates of missing data and biases. Their accuracy and efficiency outperform alternative conventional methods tested. Remarkably, EcoVAE-c can accurately reconstruct plant distributions across all genera using as little as 5% of randomly selected occurrence records. We further demonstrate the broad applicability of EcoVAE by applying it to 68 million occurrence records of butterflies and 22 million records of mammals for both genera and species (Supplementary Table 1). Additionally, our model predictions offer an unsupervised approach to assess collection completeness across different taxa at a global scale (Fig. 1b). Lastly, EcoVAE-o enables conditional predictions of species distributions based on the presence of other species (Fig. 1b), revealing ecological associations that could be validated by independent external datasets (GLoBI^34^). Our results demonstrate the unprecedented capacity and promise of deep learning methods to decode and predict biodiversity patterns at global scales.

## Results

### Overview of EcoVAE and the methodological framework

EcoVAE applies a masked approach^35^ to model the global species distributions using well-curated occurrence data based on vouchered specimen records (Fig. 1a). We applied two ranked categories for our unit of inference: genera and species. Species is the longstanding metric of inference, and genera represent coherent, morphologically similar, and often monophyletic groups of species. The later provides a practical and reasonable compromise between taxonomic detail and manageable computational demands.

The input data was converted into different resolutions, where plant presences within each grid were summarized into vectors. The richness of genera per grid varied widely, ranging from 10^1^ to 10^3^. Our model consists of an encoder that learns a low-dimensional representation of the input data and a decoder that reconstructs the presence of genera per grid. For EcoVAE-o, we randomly masked 50% of the genera presence data and the model was trained to predict these masked genera based on the remaining observed data (Fig. 1a). We did not supervise on environmental covariates, as the input co-occurrence vector serves as both input and reconstruction target when masking. This process allows the model to interpolate sparse observations and estimate the probability of unobserved plant distributions. For EcoVAE-c, bioclimatic covariates are provided as auxiliary inputs, but the training target remains the masked co-occurrence vector. EcoVAE-c is therefore able to predict presences in data sparse regions. We applied a hyperparameter grid search to select the best model configuration (Supplementary Table 2 &3, Supplementary Fig. 1&2).

### EcoVAE achieves high accuracy across regions with uneven data distribution

We evaluated the performance of EcoVAE on the plant genus dataset using a five-fold spatial block cross-validation strategy. We divided global terrestrial regions into non-overlapping 10° × 10° spatial blocks and randomly assigned them to one of five folds (Methods, Fig. 2a). In each round, we trained on spatial blocks from four folds and evaluated on the withheld (test) fold. Model accuracy was quantified using two metrics: the area under the receiver operating characteristic curve (AUROC) and the maximum True Skill Statistic (maxTSS). Both variants of EcoVAE achieved high AUROC across different folds. The mean AUROC was 0.974 (± 0.001) for EcoVAE-o and 0.973 (± 0.002) for EcoVAE-c, with corresponding average maxTSS values of 0.889 (± 0.004) and 0.887 (± 0.007). The average AUROC for each test block was largely uniform across all 10-degree blocks, demonstrating robustness despite pronounced geographic imbalances in sampling density (Fig. 2b&c). This result indicates that EcoVAE facilitates effective transferability of predictive capabilities from regions with extensive sampling to those that are sparsely sampled.

**Figure 2.**
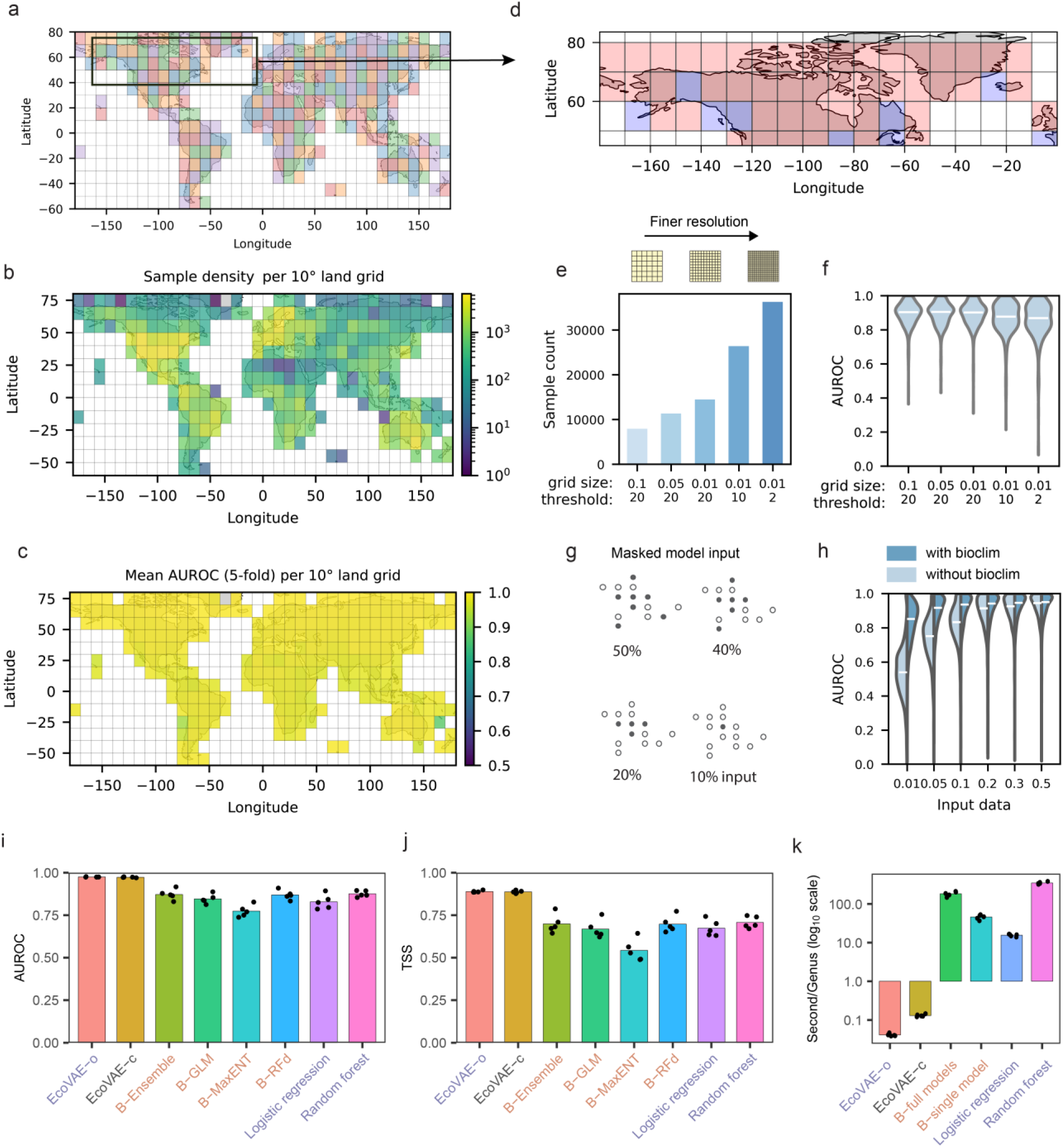
EcoVAE achieves high accuracy across regions with uneven data distribution. **a,** Training and testing blocks for each fold in spatial block cross validation. Each fold is represented by a different color. **b,** Sample density per 10° × 10° spatial block. **c,** Average AUROC for each spatial block computed on the held-out test fold. **d,** Spatial training/testing split for evaluation of model performance under sparse input. Red: training region, blue: testing region. **e,** Number of grid cells in the testing set as a function of grid size and filtering threshold. **f,** AUROC (area under the receiver operating characteristic curve) across grid sizes and filtering thresholds. **g,** Input presence records were randomly masked to simulate reduced coverage. **h,** Model performance of the test region versus the proportion of input data. Light blue: EcoVAE –o (no climatic covariates). Dark blue: EcoVAE-c (both climatic covariates and presence input). **i,** Bar plot showing AUROC values of EcoVAE-o, EcoVAE-c and SDM models in the spatial block cross-validation test. **j,** Bar plot showing TSS values of EcoVAE and SDM models. **k,** Training time per species. Models start with “B-“ are from Biomod2 package. Model names in panels i–k use the following colors: purple (presence-only input), orange (bioclimatic input), black (hybrid). Results shown are from five-fold spatial block cross-validation (n = 5).

To assess EcoVAE’s sensitivity to spatial resolutions and data filtering strategy, we conducted additional experiments focusing on the high-latitude areas of North America and the North Atlantic (Fig. 2d, including Canada, Iceland, and portions of the United States and the United Kingdom; latitude 45°–83.5°, longitude –180°–0°). We increased the spatial resolution by decreasing the grid size from 0.1° to 0.01°, maintaining the same five-fold partitioning framework. In addition, we included grids with very low presence records (grids with >= 2 genera). Such procedures increase the sample size from < 1×10⁴ to > 3×10⁴ (Fig. 2e), efficiently creating sparser input in the test grids. Across all spatial resolutions and sparsity settings, the average AUROC consistently exceeded 0.87. While no significant difference was observed with increasing spatial resolution, incorporating grids with very low occurrence records reduced predictive performance and notably increased variability (standard deviation increases from 0.07 to 0.11).

Herbarium specimen records represent a sparse sampling of actual plant distributions, and data completeness varies significantly across regions ^36,37^. To address this inherent limitation, we analyzed the impact of data sparsity on our model’s performance. We tested our model (EcoVAE-o) using only 1%, 5%, 10%, 20%, and 30% of the input genera, and evaluated its performance based on the AUROC for the remaining genera (Fig. 2g). With only 1% of the input data, the model’s performance was relatively low, with a mean AUROC of 0.54. However, increasing the input to only 5% improved the mean AUROC to 0.73. The mean AUROC further rose to 0.89 when we used 20% of the input data, close to the performance seen with 50% (Fig. 2h, AUROC of 0.92). Notably, including bioclimatic input significantly improved EcoVAE’s performance with sparse input. The mean AUROC reached 0.89 when we used only 5% of the input data and was 0.82 when 1% of the input data was fed to the model. These results demonstrate that our model enables reliable reconstruction of the full generic distribution even in data sparse regions (Fig. 2h).

### Benchmarking EcoVAE’s performance against baseline models

We subsequently compared the predictive performance of EcoVAE against widely used SDM methods. Specifically, we used four established climate-based SDM approaches: generalized linear models (GLM), down-sampled random forest (RF_d), maximum entropy modeling (MaxEnt), and an ensemble model constructed from individual models with AUROC greater than 0.8, combined through weighted averaging (Methods). We also test two machine learning models (logistic regression and random forest) that predict masked species distributions from presence-only inputs. Given the large feature space (number of genera) that significantly increases training costs, we applied factor analysis to reduce feature dimensionality prior to model fitting. We implemented the exact same five-spatial block cross validation framework for all models. Both variants of EcoVAE, with and without environmental variables, consistently outperformed all baselines with respect to both AUROC and TSS values (Fig. 2 i&j) while also demonstrating high computational efficiency (Fig. 2k).

We further compared EcoVAE with Spatial Implicit Neural Representations (SINRs), a state-of-the-art deep learning method that leverages representation learning to jointly estimate geographical range for a large number of species^38^. To further ensure a fair comparison, we applied the same dataset from the SINRs publication, i.e., presence-only data from the community science platform iNaturalist^39^. It is worth noting that the iNaturalist dataset comprises records from amateur naturalists and its sampling distribution may differ from the georeferenced vouchered specimen occurrences. We implemented the same five-fold spatial block cross-validation for both SINRs and our models. EcoVAE-o (presence-only) showed slightly lower performance comparable to SINRs. By leveraging both occurrence and bioclimatic inputs, EcoVAE-c achieved a higher AUROC than SINRs, obtaining an AUROC of 0.981 compared to 0.889 for SINRs (Supplementary Fig. 3).

### EcoVAE enables robust distribution prediction across different taxa

In addition to the plant dataset, we extended our modeling framework to other major clades with high conservation value, i.e., butterflies and mammals, and evaluated its performance using a random spatial partition approach for both genera and species.

Specifically, we selected three geographically distinct regions in North America, Europe, and Asia randomly as test areas, and applied the remaining global data for training (Fig. 3a, Supplementary Fig. 1). We calculated the predicted genera counts per grid for the test regions and compared them to the actual observations. For plants, Pearson correlation coefficients were 0.95, 0.97 and 0.97 for test regions in North America, Europe, and Asia, respectively (Fig. 3b). At the species level, our model achieved correlation coefficients of 0.95, 0.98, and 0.98 across the three regions (Supplementary Fig. 4). For the masked genera, the mean AUROC was 0.92 for North America, 0.89 for Europe, and 0.91 for Asia, which demonstrates the robust performance of our model to infer missing information from incomplete datasets (Fig. 3c). For example, *Lonicera* has a localized distribution in North America and our model correctly predicted this pattern despite this genus being masked in the input, with an overlap rate of 0.90 (Methods). Similar performance was observed for *Lamium* in Europe (0.89) and for *Rhus* in Asia (0.80), which exhibit more scattered distribution patterns (Fig. 1i). It is important to note that the three regions we selected randomly differ substantially in plant distributions, area size, and genera counts per grid (Supplementary Fig. 1, Supplementary Table 2). Nevertheless, our model performed equally well across them, which highlights the wide applicability of EcoVAE to diverse geographic contexts.

**Figure 3.**
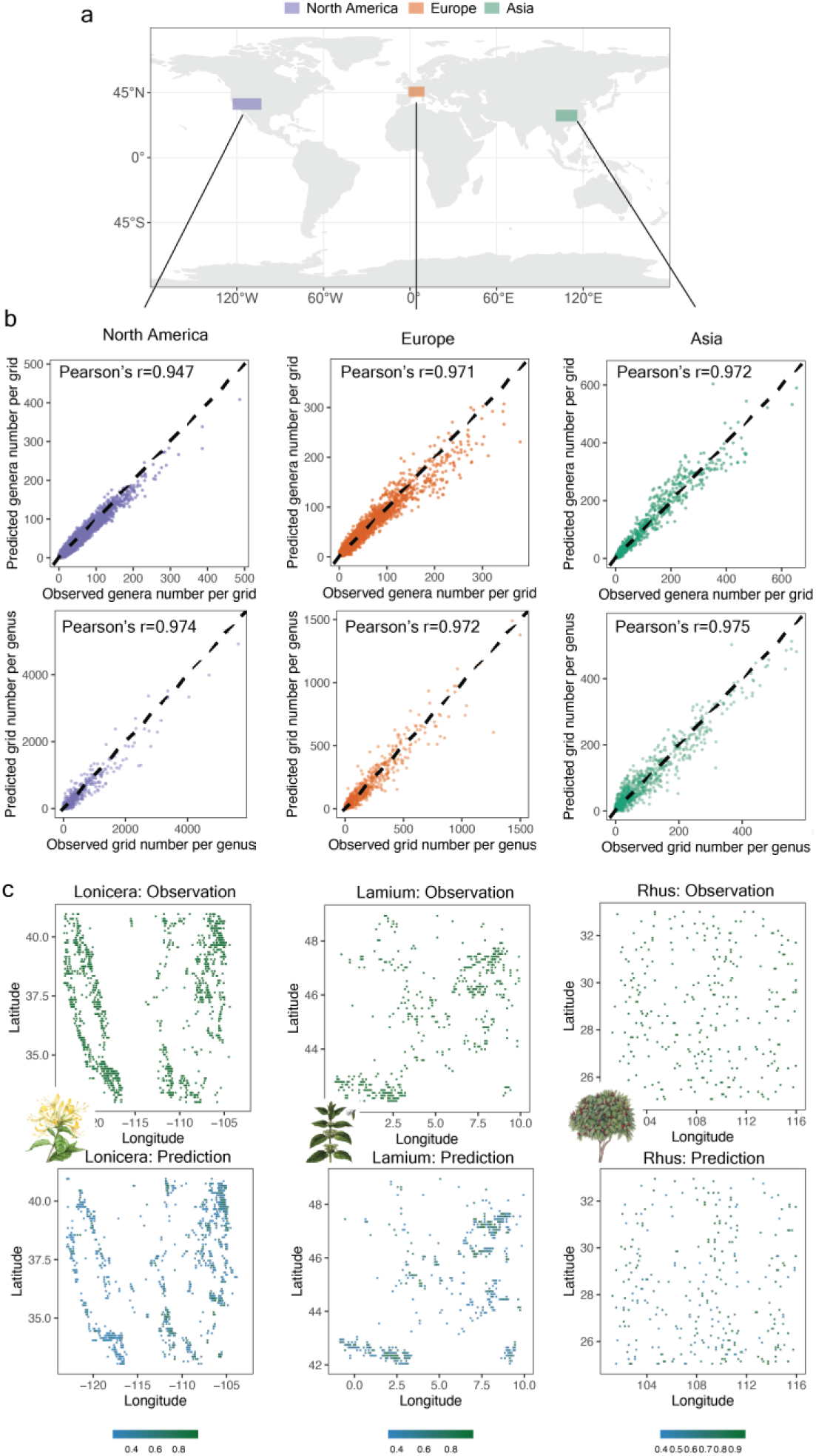
EcoVAE performs robustly in geographically distinct regions. **a,** Map showing the locations of the three testing regions. **b,** Correlation between observed genera counts per grid (or observed grid counts per genus, upper panels) and predicted genera counts per grid (or predicted grid counts per genus, lower panels) across the three testing regions, with high Pearson correlation values. The black dashed lines indicate identity lines. **c,** Comparison of the distribution of observations (upper panels) and predictions (lower panels) for randomly selected genus within each test region.

For butterflies, we found that our model achieves high accuracy in predicting the number of genera per grid, with the Pearson’s correlation coefficients of 0.92, 0.97, 0.80 for genera counts per grid for the test regions (Supplementary Fig. 5). The AUROC scores for genus-level predictions were 0.86, 0.91, and 0.77 for these regions. At the species level, the model achieved comparable results for North America and Europe, but the AUROC decreased to 0.69 for Asia, which may reflect its incompleteness of digitized vouchered occurrences at the species level (Supplementary Fig. 5). For mammals, the model performed best in the test region of North America for both genera and species (Supplementary Fig. 6). In contrast, the sparser data in Asia posed challenges for reconstructing full species distributions, which reflects the impact of uneven sampling effort. Spatial block cross-validation test confirmed our model’s robust performance across butterfly and mammal datasets (Supplementary Table 4). Overall, our results demonstrate that EcoVAE generalizes effectively across diverse taxa and geographies.

### EcoVAE predicts potential occurrences through interpolation

One important application of species modeling is interpolating occurrences where data are lacking (Fig. 4a). We hypothesize that the predictive error exhibited by EcoVAE reflects the completeness of the underlying occurrence records: if the records are incomplete, the model will struggle to reconstruct the input data effectively. We estimated the prediction error globally (Methods) and identified that our results largely reflect the known spatial biases in biodiversity sampling efforts (Supplementary Fig. 7). There is a strong correspondence between regions with high prediction error and the gaps in biodiversity monitoring^40^. This overlap is consistent with known “dark spots” of biodiversity collection estimated from expert maps and regional checklists^41,42^. For example, we observed the highest prediction errors for plants in South Asia, Southeast Asia, the Middle East, and Central Africa. South America showed higher prediction errors compared to North America (Fig. 4b). Notably, despite generally sparse records from high-latitude regions, the prediction error remained low. It could be either that the occurrence records in these areas are nearly complete, which allows the model to reflect true species distributions (Fig. 4b) more accurately, or that the species diversity is low and niches well constrained, so that few occurrences would be sufficient to capture the diversity patterns. For butterflies, the highest prediction errors were observed in South America and parts of Southeast Asia (Supplementary Fig. 8). Interestingly, central Africa exhibited low prediction errors, in contrast to the patterns observed for plants. For mammals, the prediction error was generally smaller, likely due to low generic diversity. However, regions in South America and Central Asia displayed comparatively high prediction errors, highlighting the need for further investigation in these regions.

**Figure 4.**
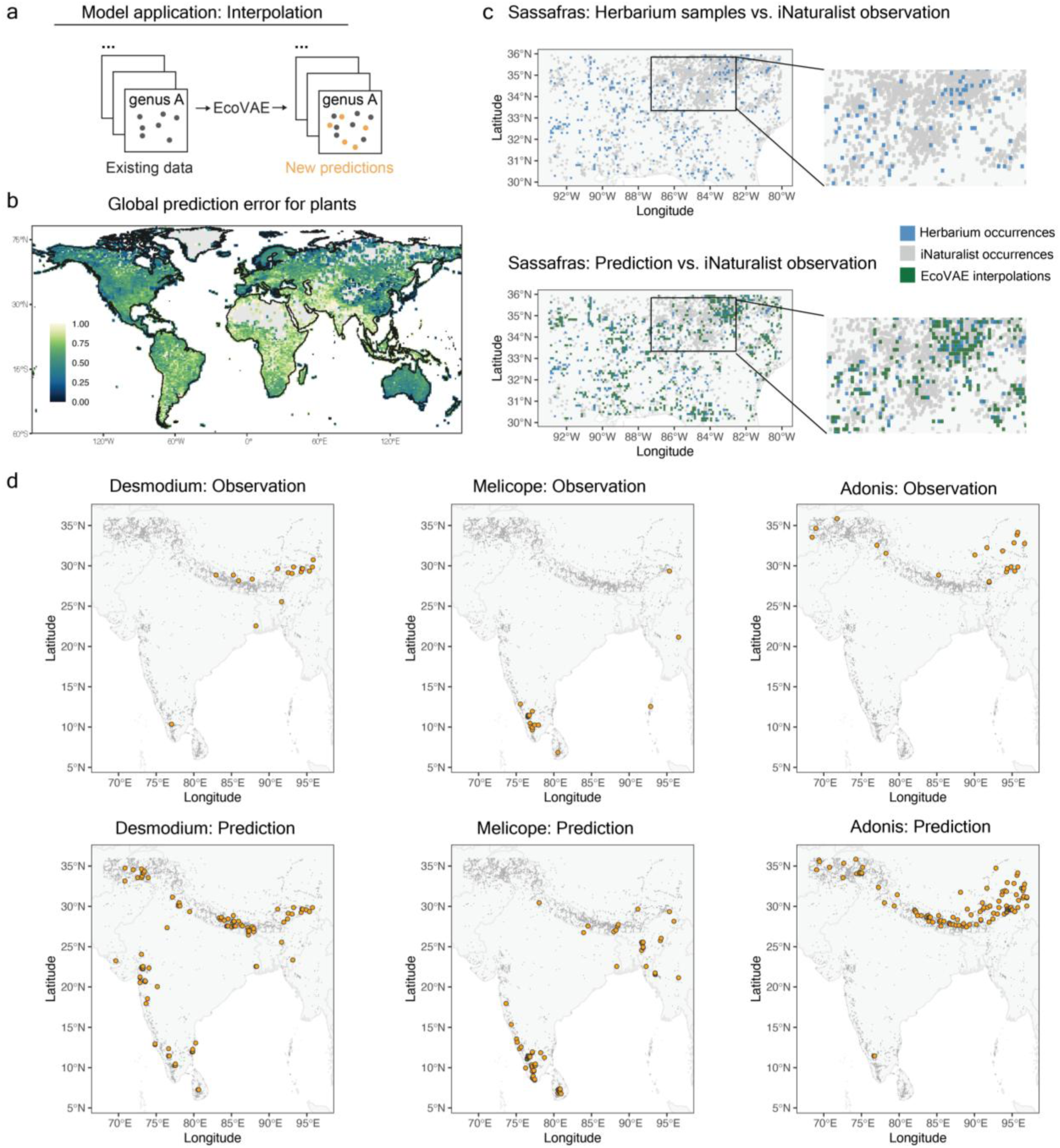
EcoVAE predicts potential occurrences through interpolation **a,** Schematic illustration of the interpolation process using EcoVAE. **b,** Global distribution of relative collection completeness, represented by the value of prediction error of EcoVAE. Darker color represents lower prediction error and higher completeness, while lighter color represents higher prediction error and lower completeness. **c,** Comparison between original herbarium specimen records and EcoVAE interpolation results of genus Sassafras in North America. **d,** Interpolation results of three example genera in South Asia. Gray dots show the distribution of all georeferenced vouchered occurrences in the study area.

We then assessed the interpolation power of EcoVAE on i.) a region in southeastern North America with relatively incomplete herbarium records but rich observational data from iNaturalist, and ii.) a region in South Asia with sparse online occurrence records of both kinds (Fig. 4a). We applied the same model structure to train a full global model based on all available plant voucher records and generated the new predictions in the two test regions (Methods). In North America, we calculated an overlap index for each genus with iNaturalist observations, defined as the ratio of predictions that are absent from input data but present in the iNaturalist data. We identified that our model performed best for genera with a moderate number of observations, while abundant data results in diminishing returns from interpolation (Supplementary Fig. 9). Using the genus *Sassafras* as an example, we identified that the new predictions largely overlap with the iNaturalist data (Fig. 4c).

For regions like South Asia, both georeferenced vouchered and observational datasets are sparser. We selected genera that showed significant expansion in our model’s new predictions. For example, the genus *Desmodium* in the Fabaceae family only have vouchered specimens in the eastern Himalayan/Nepal region in GBIF, but our model predicts its much wider distribution across western and southern India (Fig. 4d), which aligns better with field surveys and floristic investigations^43,44^. Similarly, for *Melicope* in Rutaceae, the original observations were localized in southern India, but our new predictions included occurrences in Myanmar and lowlands of Nepal, which is also confirmed by third-party observations (Fig. 4d)^45,46^. For *Adonis*, our new predictions suggested a broad distribution across the Himalayan region, consistent with various local floras and checklists describing the widespread nature of the genus from Pakistan to temperate regions in China (Fig. 4d) ^47^. These results highlight the power of our model to uncover plant distribution patterns in regions where observational data are limited.

### EcoVAE allows inference of species associations for community analysis

Another important objective of species distribution modeling is to predict community response to changing distributions of organisms within it. This is particularly relevant in invasion ecology, where it is critical to identify species vulnerable to biological invasions^22^. Our model variant, EcoVAE-o, addresses this need by generating conditional predictions: estimating the probability of presence of one (or more) species given the presence or absence of one (or more) other species.

Here, we interrogated our full global model to study genus associations based on conditional predictions. We selected a test region in Australia with abundant occurrences data and good representation of key biomes (Methods, Supplementary Fig. 10, Supplementary Table 1). We conducted *in silico* perturbation analysis, where each grid cell in the region was artificially altered by introducing a genus to areas where it was previously absent. By comparing the perturbed predictions to unperturbed models (Fig. 5a), we assessed the invasive potential of one genus on others’ distributions. We focused on statistically significant interactions for downstream analysis (Methods, Supplementary Fig. 11).

**Figure 5.**
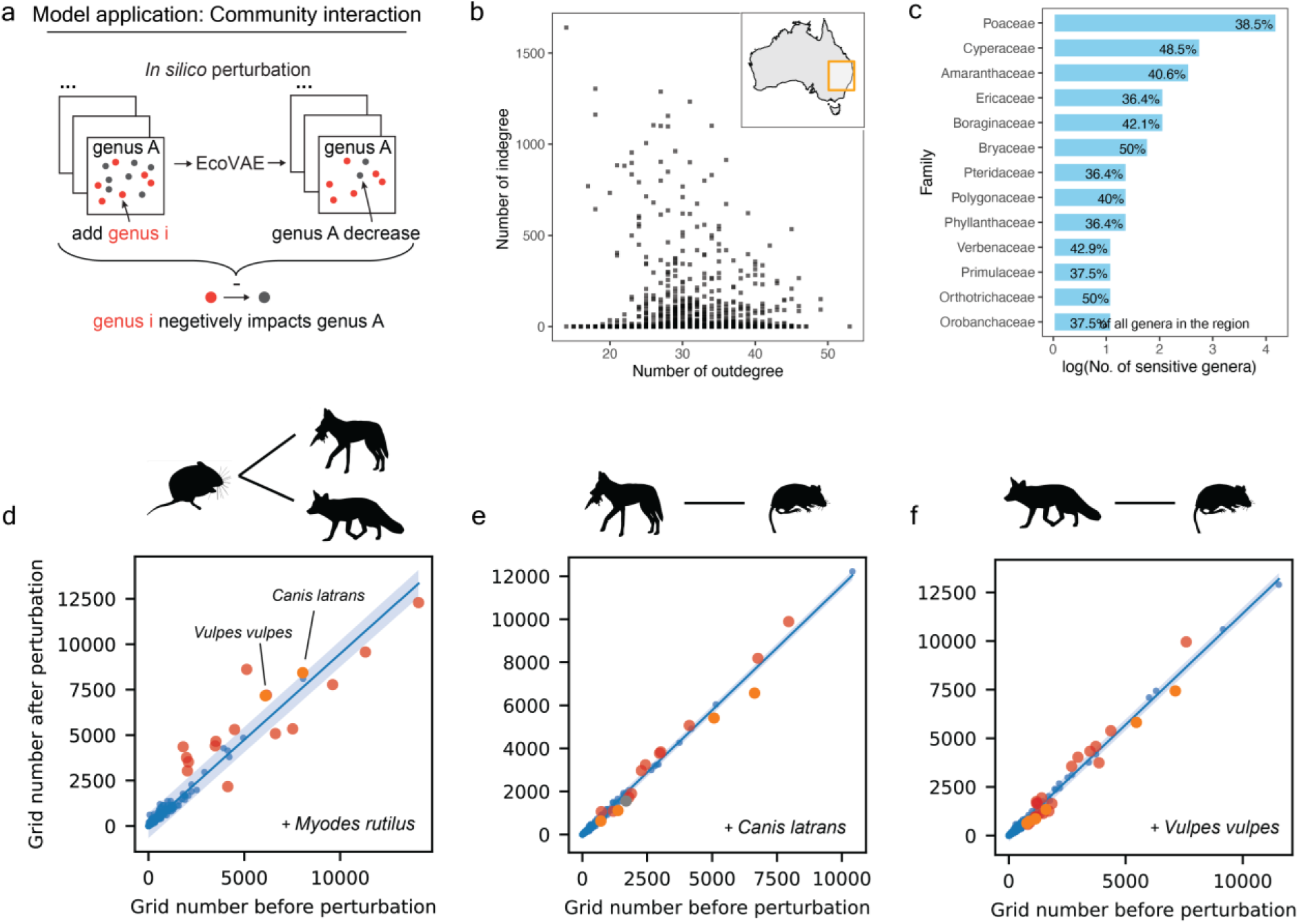
EcoVAE allows inference of species associations for community analysis a, Schematic illustration of studying community interactions using EcoVAE model. **b,** Relationship between the number of outdegree and indegree for the genus-to-genus interactions. Each dot represents a single genus. **c,** Log number of sensitive genera across the most sensitive plant families identified by EcoVAE. **d to f,** Examples of identified species associations for mammals in North America. Blue shaded area shows the 99% confidence interval of the linear regression. Each dot represents a single species. Red/orange dots represent outlier species. **d,** Perturbation by adding *Myodes rutilus*; **e,** Perturbation by adding *Canis latrans*. Orange dots denote species from *Peromyscus* genus; **f,** Perturbation by adding *Vulpes vulpes*. Orange dots denote species from *Peromyscus* genus.

Examination of our genus network revealed that genera with high out-degree (those influencing others significantly) tend to have low in-degree (being influenced by others), suggesting asymmetric interactions (Fig. 5b). Genera with broader ranges tend to interact with a larger number of genera (Supplementary Fig. 12). We revealed that certain families, including Poaceae, Cyperaceae, and Amaranthaceae, are more sensitive to disturbance in the study region (Fig. 5c). These families were significantly influenced by members from Poaceae, Asteraceae, and Fabaceae, which are globally well-represented in naturalized and invasive floras (Supplementary Fig. 13)^48^. Thus, our model reveals patterns of generic associations, providing insights into community dynamics that are generally not directly observable in the original co-occurrence records.

We further applied our model to explore interspecies association between mammals in North America. The simulated addition of *Myodes rutilus* (northern red-backed vole) increases the distribution of two predators including *Canis latrans* (coyote) and *Vulpes vulpes* (red fox) (Fig. 5d). Interestingly, these two findings align with GLoBI observations^49,50^ (Supplementary Table 5). Furthermore, simulating the addition of *Canis latrans* and *Vulpes vulpes* led to predicted changes in the distribution of various members of *Peromyscus* genus (deer mouse), the most abundant mammal in North America^51^ (Fig. 5d). While this specific pattern is not recorded in GloBI, it is consistent with known prey-predator relationships in North American small mammal communities^51,52^. All together, these results illustrate that, although statistical co-occurrence alone cannot disentangle direct from indirect effects^53^, our conditional predictions generate credible testable hypotheses about potential predator–prey links that can guide targeted field surveys or experimental follow-ups.

## Discussion

In this work, we present EcoVAE, a generative deep learning method for modeling global biodiversity distributions with high precision and speed using presence-only data, one of the most accessible data types in biodiversity science. EcoVAE provides a unified and scalable approach to evaluating global sampling efforts, interpolating distributions, and inferring interspecies associations via *in silico* perturbation experiments, offering novel tools for biodiversity exploration, monitoring, and hypothesis generation.

Our results demonstrate that species distributions can be effectively reconstructed using co-occurrence information, even with incomplete data, capturing ecological patterns often missed by traditional approaches. Adding environmental input, such as climate data, further improves model performance, making it applicable to under-sampled regions and rare species. In spatial block cross-validation tests, EcoVAE consistently outperforms traditional classifier models, and our climate-augmented variant provides additional advantages relative to the deep learning model SINRs. We propose that EcoVAE can be conceptualized as a nonlinear extension of latent factor models such as generalized linear model (GLM)-based SDM and JSDM^22^ (Supplementary Note 2). However, compared with JSDMs and other multivariate frameworks, EcoVAE is distinguished by its flexible structure for modeling multimodal inputs, its capacity to accommodate massive sample sizes, and its ability to learn complex, nonlinear dependencies between species and environments.

Although its conditional predictions are not direct measures of species interactions^53^, they nonetheless offer a foundation for applications in invasion risk assessment, and community assembly analysis^22^.

EcoVAE presents a first step towards generative modeling of co-occurrence datasets, and we acknowledge several limitations. First, our model’s performance is still constrained by the sampling bias of the occurrence record, where sampling effort is geographically uneven and certain species are more likely to be sampled or monitored. Second, EcoVAE cannot infer direct species interactions or disentangle coexistence from causal relationships^53^, since detecting interactions typically requires targeted observational or experimental evidence. Third, the nonlinearity of the encoder and decoder in EcoVAE model prevents direct estimation of regression coefficients for covariate effects on species distributions. Post hoc methods such as SHAP^54^ are needed to identify which features drive predictions and may help address potential biases.

Despite these limitations, EcoVAE complements current SDMs and JSDMs by providing a scalable framework for global analysis that can guide more targeted ecological studies. For instance, it can identify under-sampled regions or unexpected patterns, directing SDM efforts and field surveys to areas most in need of investigation. While we have demonstrated the high performance of EcoVAE on taxa such as plants, butterflies, and mammals, it can be easily extended to other taxa including birds and invertebrates, or even microbial communities. Moreover, it can be used to simulate community dynamics at macroecological scales or to test ecological hypotheses by systematically varying bioclimatic covariates or species presences, which could inform field-based approaches^15,55^. We envision that EcoVAE will advance biodiversity investigations, especially in under-sampled regions, and ultimately support global biodiversity monitoring efforts aligned with the Convention on Biological Diversity and emerging conservation frameworks such as the Kunming-Montreal Biodiversity Framwork^39^.

## Methods

### Training data

We collected georeferenced presence data based on specimen records for kingdom Plantae (plant), order Lepidoptera (butterfly and moth), and class Mammalia (mammals) from the GBIF platform (www.gbif.org, Supplementary Note 1). We used the R package “U.taxonstand”^56^ to standardized names and “CoordinateCleaner”^57^ to remove records located in the sea, on country or major area centroids, capitals, or in major biodiversity facilities. Our cleaned observation dataset included 33.8 million observations of plant, 67.6 million observations of butterfly and 21.9 million observations for mammal (Supplementary Table 2). We further used the gbifdb package (https://github.com/ropensci/gbifdb) to download coordinates of all GBIF records, which were further used to reproduce findings in Chapman et al ^58^.

The default grain size of model used in our analysis was set to 0.1’ x 0.1’. For any genus or species with more than one observation, we set the value to 1. We treat 0 as non-detection rather than confirmed ecological absence. For plant observations, we only kept genera or species that occur in over 20 grid cells and grid cells including more than five different genera/species. The finalized data contains 277,133 grids, while the three test regions include 11,292, 3,878 and 1,210 grids for North America, Europe, and Asia respectively. For butterfly and mammal observations, we only kept genera or species that occur in more than five grid cells, and grid cells with more than one genus or species.

We used the 19 bioclimatic variables from CHELSA Version 2.1^59^ at 30 arc sec (approximately 1km) per-pixel resolution for model with environmental covariates. All variables at global scale were directly downloaded from CHELSA repository (https://chelsa-climate.org/downloads/). Before fitting into the EcoVAE-c model, they were z-score normalized and clip the extreme values to –2 and 2.

### Model structure

Our core VAE architecture comprises an encoder and a decoder.

1. Encoder function: the encoder is implemented as a sequential network with two hidden layers of Gelu activated linear transformations with hidden dimension ℎ_*d*_. The encoder maps the input data (e.g., dimension: 11,555 for plant genera) into a latent space characterized by mean (μ) and log variance (*log* (σ^2^) parameters, with latent dimension *l*_*d*_.

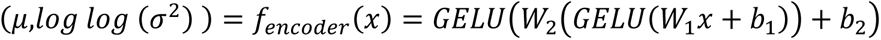

Where W1, b1, W2, b2 are the weights and biases of the two hidden layers, and GELU denotes the Gaussian Error Linear Unit activation function.
2. Reparameterization: the latent representation is obtained through a reparameterization step that ensures differentiability by sampling from a Gaussian distribution parameterized by these μ and *log* (σ^2^).

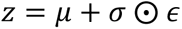

where ϵ follows normal distribution, ϵ ∼ N(0, *I*).
3. Decoder function: the decoder, mirroring the encoder’s structure, reconstructs the input data from the latent space, aiming to minimize reconstruction error. The output dimension is equivalent to the input dimension (11,555), ensuring that the reconstructed output mirrors the input feature set.

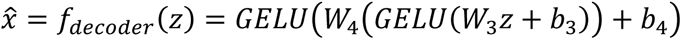

Where W3, b3, W4, b4 are the corresponding weights and biases for the decoder layers.

The model losses are measured using Mean Squared Error (MSE) or Binary Cross-Entropy (BCE).

For target *y*_*i*_ and predictions 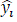 over a batch of N samples, the MSE loss can be defined as:

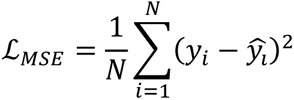

For binary target *y*_*i*_ ∈ {0, 1} and predicted probability *p*_*i*_ ∈ (0, 1), the BCE loss is:

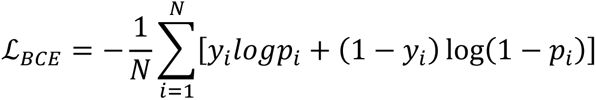

This mathematical framework enables the EcoVAE to compress high-dimensional data into a lower-dimensional latent space and subsequently reconstruct the original data with minimized reconstruction error.

### Hyper parameter scanning

To identify an efficient configuration of our EcoVAE model, we performed a targeted hyper-parameter scan covering model width, latent dimensionality, KL-divergence weighting, and loss function. We fixed the number of training epochs at ten and randomly masked 50% of all observed species/genera and used the unmasked observations to predict the distribution of the masked ones. From an exhaustive Cartesian grid, we selected combinations that spanned hidden dimensions (ℎ_*d*_) from 256 to 2,048, latent dimensions (*l*_*d*_) from 32 to 96, and BCE or MSE loss function while restricting other factors to well-performing ranges observed in preliminary trials (Supplementary Table 3).

Each model was initialized with identical weight seeds, trained on the full presence–only matrix using the Adam optimizer, and monitored for training and evaluation loss every five epochs.

### Model training and evaluation

During the model training process, we used a unique masking strategy where 50% of input data are randomly set to zero to simulate missing data scenarios. The model uses a weighted reconstruction loss function that is based on MSE or BCE. A weighting factor of 0.5 modifies the contribution of masked and unmasked genera to the reconstruction loss, providing a balanced approach to learning from both visible and obscured portions of the data. We used the Adam optimizer with a fixed learning rate of 0.001, and the models were trained for 15 epochs with a batch size of 512.

To assess model robustness while accounting for potential data leakage, we implemented a land-based spatial block cross-validation scheme. The terrestrial domain between 60° S and 80° N was first partitioned into equal 10° × 10° blocks in longitude–latitude space. Using the Natural Earth 1:110m land polygons (ne_110m_land.shp), we retained only those blocks whose outlines intersect land and randomly assigned them, under a fixed seed of 42, to one of five folds. For each fold we treated the designated land blocks as the evaluation zone and all remaining blocks as the training zone. We also dropped any test points that fell within 0.1° of an evaluation-block edge to prevent overlap between training and test areas, a step that removed roughly 4% of the evaluation records on average. The procedure yields complementary masks that guarantee strict geographic separation between training and test data while preserving the native distribution of sampling locations.

In each round, we trained the EcoVAE model on four folds and evaluated on the held-out fold. We reported area under the receiver operating characteristic curve (AUROC), Pearson and Spearman correlations (across species and across samples), and the maximum True Skill Statistic (TSS) using prevalence-matched thresholds. All metrics were computed only on masked species/genera during evaluation.

We further evaluated the model in a user-defined setting across three geographically disjoint regions with different sampling levels, including parts of Europe, western North America, and eastern Asia. We excluded data from these regions during the training phase. After model training, we conducted a masking procedure where we randomly selected 50% of all genera or species from these regions and set their corresponding observational data to zero. These masked data were then used as inputs to evaluate the model’s ability to reconstruct the observational data for the masked genera or species from the unmasked genera. For each test region, we reported the same metrics as in the spatial block cross-validation tests. In addition, for each genus or species in the test regions, we selected an equal number of top predicted grids as in the original data. We calculated the overlap rate as the fraction of predictions that had a true occurrence record either within that grid or in neighboring grids.

### Evaluate model performance on finer resolution data

We selected a test area covering parts of North America and Europe to create finer-resolution, sparser plant co-occurrence data for model evaluation. The grid cell size was reduced from 0.1° to 0.05°, 0.02°, and 0.01° (progressively finer resolutions). To further increase data sparsity, we selected grid cells with more than 2 different genera/species, instead of the original threshold of 5. Under these settings, the number of grid samples ranged from <10,000 to >30,000. For training, we fit EcoVAE on the original global dataset with grid size of 0.1° while withholding the test areas. We then tested on the withheld higher-resolution, sparser data and reported performance using AUROC.

### Evaluation of mask ratio on model performance

To assess the impact of input data completeness on model performance, we conducted a masking experiment using the global plant genus distribution dataset. As in the benchmarking experiments, we withheld three geographic regions from the training set and used them exclusively for evaluation. Within each test region, we randomly retained a fixed proportion of the observed genera—specifically 0.01, 0.05, 0.1, 0.2, 0.3, and 0.5—and masked the remainder. We then evaluated EcoVAE’s ability to reconstruct the masked genus-level distribution based on the partial input. Two model variants were tested: a presence-only model (EcoVAE-o) and a model incorporating both environmental covariates and presence data (EcoVAE-c).

### Single-species distribution models based on presence-only data

To evaluate the time efficiency of our EcoVAE model in processing large-scale input data, we conducted a series of benchmarking experiments in comparison with two popular SDM methods, i.e., random forest and logistic regression^43^. We tested the model’s performance under varying input dimensions by randomly selecting genera from the input genus presence matrix. For these experiments, we used a masking strategy where 80% of the input columns were masked to simulate missing data. A binary mask was generated using a probability threshold proportional to the desired masking percentage, ensuring that columns in the input data were set to zero with the defined probability. We used the training and test data split as previously described. A random genus with presence in at least five grids was randomly chosen as the prediction target: the model should predict the presence of this genus across all grids in the training and test data based on the masked input matrix. We trained the random forest classifier using the *sklearn.ensemble.RandomForestClassifier* function with default parameters. We trained the logistic regression model with the *sklearn.linear_model. LogisticRegression* function with the following parameters: max_iter=1000, solver=’saga’, penalty=’l1’. Time measurements were recorded from the start of data preparation to the completion of the prediction phase. The time consumption was calculated over 10 iterations to benchmark the models’ time efficiency. To further reduce input dimension for the machine learning models, we applied factor analysis to the presence matrix using s*klearn.decomposition.FactorAnalysis* function with default parameters. We reduced the feature dimension to 100 or 200, then trained the models and evaluated runtime and performance as described above.

### Single-species distribution models based on bioclimatic variables

We evaluated commonly used SDMs with the R package biomod2 (v4.3)^60,61^, benchmarking three widely used algorithms for performance: generalized linear models (GLMs), MaxENT, and a down sampled, random forest ensemble (RFd)^62^. We also conducted an ensemble modeling method, which was constructed by weighting individual model predictions using AUC-derived weights, retaining only models with AUC >0.8^63^. We randomly selected 1000 plant genera and species for SDM from the baseline presence dataset generated for EcoVAE, calculated the occurrences in each spatial block following the same cross-validation framework. We further filtered taxa to ensure adequate information for single-species SDMs using three criteria: total occurrence records (>50), minimum occurrences in the testing set (>5), and representation in any given spatial block constituting <20% of all presence records. These filters ensured robust model training and reduced spatial autocorrelation bias. For each taxon, Bioclim variables were clipped to a +10° bounding box around all training occurrences. Within this cropped extent, 5,000 pseudo-absences were sampled randomly. To address collinearity, we also used a flexible variable selection strategy based on sample size: for species with <200 occurrences, a reduced, uncorrelated subset (Pearson’s |r| < 0.7) of predictors was selected from the most informative variables, with the number of predictors limited to either ⌊n/10⌋ or the smallest subset meeting the correlation criteria. Species with ≥ 200 occurrences used all 19 variables ^64^. Model hyperparameters followed the optimized “bigboss” settings recommended in the biomod2 documentation for large-scale applications. Evaluation metrics included: area under the receiver operating characteristic curve (AUC), the true skill statistic (TSS), and the continuous Boyce index. All modeling was run on the Harvard Cannon cluster using 8 cores, and processing time was recorded for each modeling run. In addition, we trained logistic regression and random forests models with *scikit-learn* package, using bioclimatic variables to predict species occurrence. The protocol mirrored the presence-only experiments, except that the inputs were the bioclimatic predictors.

### Benchmarking Spatial Implicit Neural Representations (SINRs)

We benchmarked the performance of EcoVAE against SINRs^38^, a state-ot-the-art deep learning method for predicting species distributions based on climatic covariates. We followed the original paper’s data-loading and preprocessing pipeline for the iNaturalist dataset and used the default training parameters from the function *setup.get_default_params_train*. We trained SINRs with the ℒ_*A*N−*full*_loss since it tends to perform better than the others^38^. The only modification was the evaluation protocol and we implemented the same five-fold spatial block cross-validation used for EcoVAE. Specially, the terrestrial domain between 60° S and 80° N was divided into 10° × 10° blocks. Using the Natural Earth 1:110m land polygons (ne_110m_land.shp), we retained only those blocks whose outlines intersect land and randomly assigned them, to one of five folds. For each fold we treated the designated land blocks as the evaluation zone and all remaining blocks as the training zone. We also dropped any test points that fell within 0.1° of an evaluation-block edge to prevent overlap between training and test areas. The model performance was measured using AUROC, AUPRC and maxTSS. We further aggregated the same iNaturalist dataset into 0.1° × 0.1° grids and evaluated EcoVAE under the same spatial block cross-validation protocol.

## Model application

### -Data interpolation

We trained the EcoVAE model as previously described on all available occurrence data for interpolating unobserved plant distribution (full global model). To evaluate the model’s prediction error, we used the unmasked global data as input and applied a threshold to binarize the output genus presence matrix, ensuring that the total number of occurrences was doubled. For each grid, we then calculated the ratio of observed genera not represented in the output matrix, which we defined as the prediction error. Based on the global distribution of prediction error, we selected two regions to evaluate the performance of data interpolation, i.e., North America and South Asia. For these regions, we used a similar strategy to generate the binarized output for downstream analysis.

For the North America region, we used observation data collected from iNaturalist to verify the accuracy of model prediction instead of traditional data splitting method. The ratio between iNaturalist observational data and vouchered specimen data is 7:1 and we would expect that iNaturalist data have a better geographic coverage than herbarium specimens for many species. We calculated overlapping rate between predicted species occurrences and actual observations to quantify the performance of our prediction. For each species, we first extracted the observed and predicted occurrences based on the genus index. For both datasets, we retained only the presence points. The observed points were buffered by 0.1 degrees to account for geographic uncertainties. We then converted the presence data into spatial objects using the “sf” package in R^65^. Overlap between predicted occurrences and observed occurrences, as well as overlap between predicted and original input occurrences, was calculated using spatial intersections (“st_intersects2019;). The key metric, the overlapping rate, was calculated by dividing the number of predicted points that overlapped with observed points but not with original input points by the total number of predicted points. This rate reflects the proportion of new predicted occurrences that align with observed data but were not part of the input data, providing a measure of prediction accuracy and novelty. We also compared the predicted overlapping rate with the overlapping rate calculated between the same number of randomly generated points in the study area and observations.

For the South Asia region, we assessed the occurrence of each genus both before and after data interpolation. We focused on genera that initially occurred in more than 5 grids and whose distribution region has expanded most for downstream analysis. Due to the lack of georeferenced observational data in this region, we compared our prediction with the distribution described in plant atlas and related literature.

### -Simulation of genera associations

To simulate the impact of a specific genus i on all other genera within a targeted region, we initially identified all grid cells lacking genus i. The observational data from these grids were utilized as input for the model, and the corresponding reconstructed data served as the background dataset (x_background). Then we introduced observations of genus i into these grids and generated perturbed model outputs (x_perturb). By comparing the plant distributions between x_background and x_perturb, we were able to identify genera that exhibited significant changes, thereby quantifying the ecological influence of genus i on the plant community dynamics within the region.

To assess species associations after species additions, we first fit linear regression models to compare grid numbers of all genera before and after addition of a specific genus i. Specifically, we used the ‘lm’ function in R to model the relationship between grid numbers before and after addition of genus i. For each model, we calculated 99.99% confidence intervals using the ‘predict’ function with the interval parameter of “prediction” and level parameter of 0.9999. We defined the significant interaction between genus i and j if the predicted grid number for genus j falls outside the bounds of the confidence intervals after addition of genus i. In such circumstances, we identified j as an “outlier” and defined it as a “sensitive genus”. We used the same protocol to identify species associations for mammals in North America, and the only difference is that we calculated 99% confidence intervals. Z-scores were calculated for each genus by normalizing the residuals, computed as the difference between actual and predicted values. We then performed frequency analysis of the impactful and vulnerable genera based on all significant interactions. To further explore genus interactions at the family level, we utilized the “plantlist” package ^66^ to classify genera into families and analyzed the proportion of sensitive genera within each family. We selected the most sensitive family based on the following criteria: it includes more than 5 genera and at least 35% of the genera are classified as “sensitive” (significantly impacted by at least one other genus).

## Code Availability

Our trained model and codes are available from GitHub: https://github.com/EcoVAE/EcoVAE

## Author contributions

YY and SB conceptualized the study, conceived EcoVAE, collected and analyzed the data. YY, JY, and SB wrote the manuscript with key contributions from CCD. All authors approved the manuscript.

## Acknowledgements

YY was supported by Harvard University. CCD was supported by Harvard University, LVMH Research, Dior Science, and National Science Foundation grants DEB-1355064 and DEB-0544039. We thank Sound Solution for valuable suggestions.

