## Supplementary Information for "A generative deep learning method for global species distribution prediction"

**Supplementary Note 1.** Plant occurrence records since 1900 were downloaded from GBIF in April 2024 using r package "rgbif". Due to the download size limitation of the package, the global plant distribution data was divided into six smaller dataset based on latitude and longitude and then combined. Each dataset was inquired using the inquiry as followed:

```
occ_download(type="and",
                    #pred_not("taxonKey", 212),
                    pred("kingdomKey", 6),
                    pred("hasGeospatialIssue", FALSE),
                    pred("hasCoordinate", TRUE),
                    pred("occurrenceStatus","PRESENT"),
                    #pred_gte("year", 1900),
                    #pred("basisOfRecord","Preserved specimen"),
                    pred_within(polygons[[4]]),
                    pred_or(
                    pred_not(pred_in("establishmentMeans",c("MANAGED","INTRODUCED"))),
                      pred_isnull("establishmentMeans"),
format = "SIMPLE_CSV")
```

The doi of each datasets are: DOI10.15468/dl.cj23sr, DOI10.15468/dl.fhje4h, DOI10.15468/dl.s59um3, DOI10.15468/dl.6sp4gd, DOI10.15468/dl.qq5ngz, DOI10.15468/dl.vt4mzs

Mammal and butterfly occurrence records were downloaded from GBIF through the following urls: https://doi.org/10.15468/dl.ed3xvf, https://doi.org/10.15468/dl.utzfa3

### Supplementary Note 2. EcoVAE as a Nonlinear Latent Factor Model

We focused on our discussion on EcoVAE-o, which is trained with the presence–only data $Y \in \{0,1\}^{N \times S}$ where $S$ is the number of species and $N$ the number of locations. Each element $y_{ij} = 1$ if species $j$ is present at location $i$. We treat 0 as non-detection rather than confirmed ecological absence. By randomly masking part of observations, EcoVAE directly learns the conditional probability of co-occurrence $P[\{y_{ij}\}_{j \notin A} | \{y_{ij}\}_{j \in A}]$, i.e., at a given location $i$, if we observe a a subset of $y_{ij}, j \in A$, how likely the rest of the species occur in the same location. In this section, we mathematically show how EcoVAE can be treated as a non-trivial extension to the existing species distribution model (SDM).

#### Species distribution models

Generically, a species distribution model aims to predict the distribution or abundance of multiple species $y_i$ at each location $i$. One fundamental assumption is that the co-occurrence data observed is a result of both environments and species-species interactions. To model such dependencies, it's usually assumed that these factors can be captured by a list of mathematically well-defined features, denoted as $X_i$ and $\Omega_i$ as environmental covariates and species-species interaction covariates. Specifically, such a model tries to learn the conditional probability:

$$P(y_i | X_i, \Omega_i).$$

However, while $X_i$ can usually be pre-defined as one can select a list of environmental covariates, $\Omega_i$ is difficult to evaluate, and is often regarded as what one aims to infer from the model instead of a set of covariates as a part of the model's input. In fact, there is a debate whether introducing $\Omega_i$ can alter the estimation of species' fundamental niches. It has been shown that it cannot, in the setting of the species distribution model (SDM) and joint species distribution model (JSDM)[1]. Here we briefly review the proof from Poggiato et al[1].

In both SDM and JSDM, the presence of species $Y$ is predicted by a logistic regression model:

$$Y = I(Z > 0), \text{eq.(1.1)}$$
$$Z = X \cdot B^\top + \epsilon, \text{eq.(1.2)}$$

where $I(\cdot)$ is the indicator function, $X \in R^{N \times K}$ are the $K$ measured environmental covariates at each location, $B \in R^{S \times K}$ is the regression coefficient matrix. The major difference between SDM and JSDM is that whether each species is modelled independently or not. In the SDM, $\epsilon_i \sim N_S(0, I)$, and in JSDM, $\epsilon_i \sim N_S(0, R)$ where $R$ is a correlation matrix which describes the residual correlation among taxa and reflects species co-occurrence patterns not explained by the selected environmental covariates.

In other words, the difference is that SDMs assume independent residuals, while JSDMs allow for correlations between them.

Many JSDMs use latent factors to reduce the dimension of $R$, e.g., to extend eq.(1.2) to:

$Z = X \cdot B^{\top} + \Omega\Lambda^{\top} + E$, eq.(1.3)

Where $\Omega \in R^{N\times L}$ is the latent factor matrix of $L$ features on $N$ locations, and $\Lambda \in R^{S\times L}$ is another regression coefficient. Due to lack of prior knowledge on how species interaction contributes to the observed distribution, latent factors are also assumed to be Gaussian, i.e. $\Omega_{il} \sim N(0,1)$. Therefore, essentially, SDM and JSDM (including the latent factor model eq.(1.3)), are equivalent to the following logistic regression model:

$Y = I(Z > 0)$,
$Z = X \cdot B^{\top} + error$,

where the error is Gaussian. The maximum likelihood estimator (MLE) of all such models, are the same regardless of the covariance matrix of the error term:

$\hat{B} = (X^{\top}X)^{-1}X^{\top}Y$.

Therefore adding $\Omega$ doesn't change the estimation of $B$, in other words, the response of $Y$ to the environmental covariates $X$ remains the same when introducing covariances between the errors or the latent factors $\Omega$. The same conclusion can also be obtained from a Bayesian perspective[2]. This result is not surprising as Gaussian noise doesn't provide any information of the effect of other species, so the estimator should not change. In the next subsection, we discuss whether it's possible to modify the SDM in the way that $\Omega$ can be inferred from the observed data.

**Variational SDM infers the latent factors from observed co-occurrence data**
As described above, the major reason why introducing $\Omega$ cannot help is because $\Omega$ can be merged into the error term, since it consists of independent and identically distributed (i.i.d.) Gaussian noise. It's then naturally to ask whether it is possible to infer the latent factor $\Omega$ itself from the co-occurrence data. In this subsection, we propose a variational inference framework for predicting the occurrence of unobserved group of species based on observed co-occurrence data, i.e., $P[\{y_{ij}\}_{j\notin A}|\{y_{ij}\}_{j\in A}]$, tentatively termed as variational SDM. For mathematical simplicity, we consider a simpler case where both latent factor $\Omega$ and environmental covariates $X$ are inferred from the co-occurrence data.

In most of the cases including the case considered in this manuscript, both $K$ (number of environmental covariates) and $L$ (number of latent factors) are much smaller than the number of species $S$, so we assume that $K + L < S$, Eq. (1.3) can then be rewritten as:

$Z = [X\Omega] \cdot [B\Lambda]^{\top} + E = \tilde{X} \cdot \tilde{B}^{\top} + E$, eq.(2.1)

where $\tilde{B} \in R^{S\times(K+L)}$, because $K + L < S$, it's reasonable to assume $\tilde{B}$ has linearly independent columns, so the right inverse of $\tilde{B}^{\top}$ exists: $\tilde{B}^{+} = \tilde{B}(\tilde{B}^{\top}\tilde{B})^{-1}$. Multiply the right inverse on both sides of Eq. (3), we have an "inverse" latent factor model:

$\tilde{X} = Z\tilde{B}^{+} + \tilde{E}$, eq.(2.2)

This equation shows that, in principle, we can infer the latent factors (along with environmental covariates) from the co-occurrence data itself through regression, and use the inferred latent factors to further predict the co-occurrence of the unobserved group of species. This approach is fundamentally different from JSDM because the latent factors are no longer i.i.d. Gaussians but directly inferred from the co-occurrence data which brings the information of species interactions and may enhance the prediction power of the model. However, this approach is difficult or even impractical to implement in the setting of Eq. (2.1-2.2), mainly because it is hard to optimise $\tilde{B}(\tilde{B}^{\top}\tilde{B})^{-1}$ directly. More importantly, the limited expressiveness of linear models makes it difficult for capturing the complex interactions between environmental factors and species dynamics. In the next subsection we show that EcoVAE solves both problems by leveraging the expressiveness of neural network and gradient-based optimization approach.

**EcoVAE is a nonlinear extension of the latent factor model**

In this subsection we briefly review Variational Autoencoder (VAE) and show it's a nonlinear extension of the variational inference model described in the previous subsection.

In a variational autoeocoder[3], the input data, e.g., $Y_A = \{y_j\}_{j\in A}$ is first encoded into a lower dimensional latent space through a nonlinear function represented by a neural network $Encoder(Y_A; \theta_E)$, where the $\theta_E$ denotes the parameters in the encoder function. Specifically, in EcoVAE, the encoder is:

$Encoder(Y_A; \theta_E) = GeLU(GeLU(Y_A \cdot W_1 + b_1) \cdot W_2 + b_2, \theta_E = \{W_1, W_2, b_1, b_2\}$,

where $GeLU$, the Gaussian Error Linear Unit[4] serves as the activation function.

The latent space variables $X$, similar to Eq. (2.2), is assumed to be Gaussian, with mean and variance depending on $Y_A$. So practically, the encoder gives the mean and variance of $X$:

$\mu_{Y_A}, \sigma^2_{Y_A} = Encoder(Y_A; \theta_E)$,

The new latent variable $\tilde{X}$ is then sampled by:

$$\tilde{X} = \mu_{Y_A} + \sigma_{Y_A} \cdot \varepsilon$$

where $\varepsilon \sim N(0,1)$.

Notice that here $\tilde{X}$ is still in the latent space but not in the dimension of $S$ (which is different from Eq. (2.2)), we need another neural network, i.e., the decoder, to transform it into our predicted occurrence data):

$\tilde{Y} = Decoder(\tilde{X}; \theta_D)$,

where the decoder in EcoVAE uses the same structure as the Encoder. The model is then trained in the way that the re-generated data $\tilde{Y}$ follows the same (or as close as possible) distribution of the original data $Y_A$, by optimizing $\theta_E$ and $\theta_D$ with the Evidence Lower Bound (ELBO) loss[3,5].

EcoVAE is a nonlinear version of the variational inference framework proposed in the previous subsection. To see this, we compare both algorithms:

| | Step | variational SDM | EcoVAE |
|---|---|---|---|
| 1 | Inferring the distribution of latent factors from co-occurrence data $Y_A$ | $Z = logit(Y_A), \mu_A = Z\tilde{B}^{+}$, $\sigma^2 = [\tilde{E}\tilde{E}^{\top}]$ is fixed | $\mu_{Y_A}, \sigma^2_{Y_A} = Encoder(Y_A; \theta_E)$ |
| 2 | Sampling latent variables $\tilde{X}$ | sampling $\tilde{X}$ from $N(\mu_A, \sigma^2)$ | sampling $\tilde{X}$ from $N(\mu_{Y_A}, \sigma^2_{Y_A})$ |
| 3 | Compute the co-occurrence $\tilde{Y}$ of unobserved species | $\tilde{Y} = I(\tilde{X} \cdot \tilde{B}^{\top} + E > 0)$ | $\tilde{Y} = Decoder(\tilde{X}; \theta_D)$ |

We can see that both algorithms have the same steps. The main difference is that the EcoVAE uses nonlinear functions encoded by neural networks, while variational SDM uses linear functions. Therefore, EcoVAE can be regarded as a nonlinear extension of the variational SDM. Compared to SDM or JSDM we summarized above, it learns the low-dimensional latent factors directly from the co-occurrence data, instead of merely sampling from univariate Gaussian distributions. Compared to the variational SDM, it leverages the expressiveness of neural networks which is more capable of capturing the highly nonlinear interactions between environments and species in complex ecological systems. Furthermore, the optimization can be easily implemented with stochastic gradient descent (SGD) algorithm.

EcoVAE reconstructs masked species to predict unobserved distributions from the observed ones, enabling accurate estimation of probability of species co-occurrences. Our approach offers conditional predictions (inference of the probabilities for one or more species, given the presence of others), without asserting explicit species interactions. We emphasize that conditional dependency learnt from large-scale co-occurrence data should not be interpreted as direct ecological interactions, as it has been thoroughly discussed in Blanchet et al. Echoing Poggiato et al., we argue that this conditional prediction capacity could be exploited for diverse ecological applications such as probing the probability for species invasion and reintroduction analysis.

**Supplementary Table 1. Statistics for vouchered records and EcoVAE models.**

| Taxa | Plantae | | Lepidoptera | | Mammalia | |
|---|---|---|---|---|---|---|
| | **genus** | **species** | **genus** | **species** | **genus** | **species** |
| **species/genus number** | 11,555 | 127,281 | 11,105 | 47,485 | 1,270 | 4,872 |
| **records** | 33.82M | 33.82M | 67.62M | 66.20M | 21.53M | 20.92M |
| **model parameter number** | 6.89M | 65.45M | 5.85M | 24.52M | 0.81M | 2.66M |

**Supplementary Table 2. The geographic range of the regions selected in the study.**

| | Region | Latitude | Longitude |
|---|---|---|---|
| **Model evaluation** | North America | 33 to 41 | -123 to -103 |
| | Europe | 42 to 49 | -1 to 10 |
| | Asia | 25 to 33 | 101 to 116 |
| **Interpolation** | North America | 30 to 36 | -93 to -80 |
| | South Asia | 6 to 36 | 68 to 97 |
| **Species interaction** | Australia | -35 to -25 | 144 to 154 |
| | North America | 15 to 60 | -110 to -75 |

**Supplementary Table 3. Hyperparameter scan for EcoVAE under 50% random masking.** We reported AUROC and maxTSS for three held-out test regions (EU, NA, AS) after 10 training epochs of the EcoVAE-o model. For each test region, the best value is shown in bold.

| Hidden dim | Latent dim | Kl weight | Loss function | Region 1 (EU) | | Region 2 (NA) | | Region 3 (AS) | |
|---|---|---|---|---|---|---|---|---|---|
| | | | | AUROC | maxTSS | AUROC | maxTSS | AUROC | maxTSS |
| 256 | 32 | 0 | mse | 0.834 | 0.638 | 0.867 | 0.699 | 0.823 | 0.653 |
| 256 | 32 | 0 | bce | 0.884 | 0.702 | 0.913 | 0.771 | 0.905 | 0.754 |
| 256 | 32 | 0.1 | mse | 0.715 | 0.500 | 0.765 | 0.568 | 0.749 | 0.557 |
| 256 | 32 | 0.1 | bce | 0.876 | 0.692 | 0.919 | 0.780 | 0.899 | 0.746 |
| 256 | 96 | 0 | bce | 0.877 | 0.692 | 0.920 | 0.780 | 0.912 | 0.764 |
| 256 | 96 | 0.1 | mse | 0.784 | 0.562 | 0.813 | 0.615 | 0.807 | 0.625 |
| 256 | 96 | 0.1 | bce | 0.876 | 0.685 | 0.910 | 0.765 | 0.909 | 0.759 |
| 768 | 32 | 0 | mse | 0.867 | 0.675 | 0.881 | 0.723 | 0.872 | 0.708 |
| 768 | 32 | 0 | bce | 0.884 | 0.706 | 0.921 | 0.786 | 0.910 | 0.762 |
| 768 | 32 | 0.1 | mse | 0.819 | 0.617 | 0.810 | 0.629 | 0.840 | 0.674 |
| 768 | 32 | 0.1 | bce | 0.878 | 0.695 | 0.916 | 0.778 | 0.912 | 0.766 |
| 768 | 96 | 0 | bce | 0.879 | 0.698 | 0.920 | 0.782 | 0.912 | 0.769 |
| 768 | 96 | 0.1 | mse | 0.832 | 0.635 | 0.845 | 0.676 | 0.839 | 0.671 |
| 768 | 96 | 0.1 | bce | 0.874 | 0.693 | 0.923 | 0.787 | 0.904 | 0.754 |
| 2048 | 32 | 0 | bce | **0.887** | **0.708** | 0.921 | 0.785 | 0.912 | 0.768 |
| 2048 | 32 | 0.1 | bce | 0.881 | 0.700 | **0.924** | **0.789** | 0.906 | 0.758 |
| 2048 | 96 | 0 | bce | 0.881 | 0.697 | 0.924 | 0.787 | **0.913** | **0.769** |
| 2048 | 96 | 0.1 | bce | 0.880 | 0.691 | 0.919 | 0.782 | 0.912 | 0.767 |

**Supplementary Table 4. EcoVAE's performance across different taxa.** Metrics (mean ± s.d.) including AUROC, AUPRC, and maxTSS are reported under presence-only ("occur") and combined presence+climate ("occur+clim") inputs.

| Dataset | Input | Parameter number | AUROC | AUPRC | maxTSS |
|---|---|---|---|---|---|
| Plantae (genus) | occur | 128.6M | 0.975 ± 0.001 | 0.245 ± 0.009 | 0.889± 0.006 |
| Plantae(genus) | occur+clim | 128.9M | 0.973± 0.002 | 0.218± 0.012 | 0.887± 0.008 |
| Mammalia (species) | occur | 23.0M | 0.927± 0.004 | 0.149± 0.016 | 0.828± 0.011 |
| Mammalia (species) | occur+ clim | 23.1M | 0.950± 0.005 | 0.162± 0.014 | 0.884± 0.010 |
| Lepidoptera (species) | occur | 154.7M | 0.957± 0.010 | 0.193± 0.008 | 0.894± 0.016 |
| Lepidoptera (species) | occur+ clim | 154.8M | 0.971± 0.004 | 0.191± 0.010 | 0.920± 0.008 |

**Supplementary Table 5. Examples of species associations validated using the GloBI dataset.**

| source_taxon_name | source_taxon_path_ids | interaction_type | target_taxon_name | target_taxon_path_ids | study_citation |
|---|---|---|---|---|---|
| Myodes rutilus | GBIF:5706762 | interactsWith | Canis latrans | GBIF:5219153 | A. Thessen. 2014. Species associations extracted from EOL text data objects via text mining. Accessed at <https://github.com/EOL/pseudonitzchia/archive/e5838965a186fba4b7215cd0d179c4526773bad5.zip> on 13 Sep 2025. |
| Myodes rutilus | GBIF:5706762 | eatenBy | Vulpes | GBIF:5219234 | Wilson, D.E. & Mittermeier, R.A. eds. (2009). Handbook of the Mammals of the World. Vol. 1. Carnivores. Lynx Edicions, Barcelona. |
| Myodes rutilus | GBIF:5706762 | eatenBy | Vulpes | GBIF:5219234 | Wilson, D.E. & Mittermeier, R.A. eds. (2009). Handbook of the Mammals of the World. Vol. 1. Carnivores. Lynx Edicions, Barcelona. |
| Myodes rutilus | GBIF:5706762 | eatenBy | Vulpes vulpes | GBIF:5219243 | Wilson, D.E. & Mittermeier, R.A. eds. (2009). Handbook of the Mammals of the World. Vol. 1. Carnivores. Lynx Edicions, Barcelona. |
| Myodes rutilus | GBIF:5706762 | eatenBy | Vulpes vulpes | GBIF:5219243 | Wilson, D.E. & Mittermeier, R.A. eds. (2009). Handbook of the Mammals of the World. Vol. 1. Carnivores. Lynx Edicions, Barcelona. |
| Myodes rutilus | GBIF:5706762 | interactsWith | Vulpes vulpes | GBIF:5219243 | A. Thessen. 2014. Species associations extracted from EOL text data objects via text mining. Accessed at <https://github.com/EOL/pseudonitzchia/archive/e5838965a186fba4b7215cd0d179c4526773bad5.zip> on 13 Sep 2025. |

**Supplementary Figure 1**. **The distribution of terrestrial plant genus number per grid. a,** all study grids; **b-d,** grids in three testing regions.

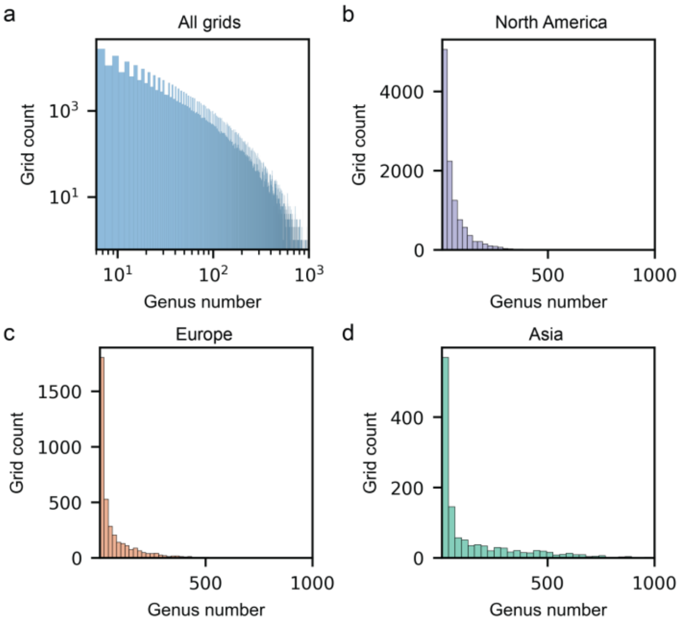

**Supplementary Figure 2. Effect of batch size on model performance. a,** Model performance (AUROC) across three test regions as a function of batch size. The model was trained on global plant distributions at the genus level. Blue: EU; Orange: NA; Green: Asia. **b,** Model training time versus batch size.

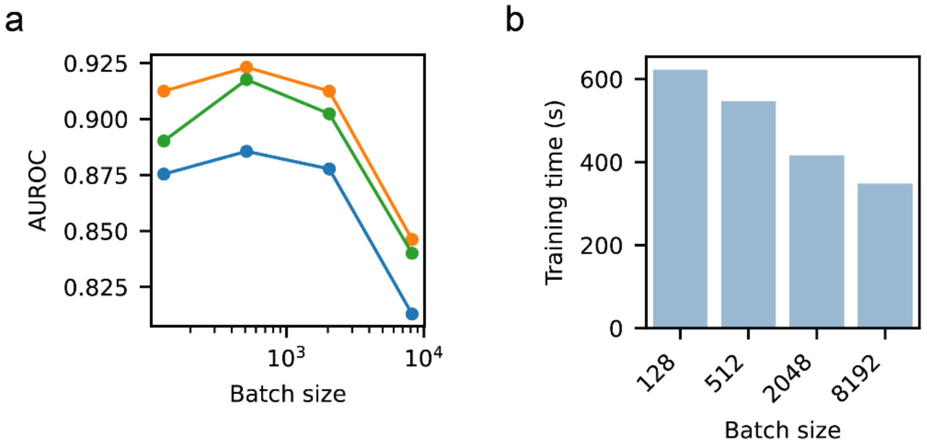

**Supplementary Figure 3. Performance of SINRs and EcoVAE on the iNaturalist dataset. a,** Sample density of the iNaturalist dataset. **b,** AUROC for EcoVAE models compared with SINRs under spatial block cross-validation tests. Each dot represents result from one fold (n = 5).

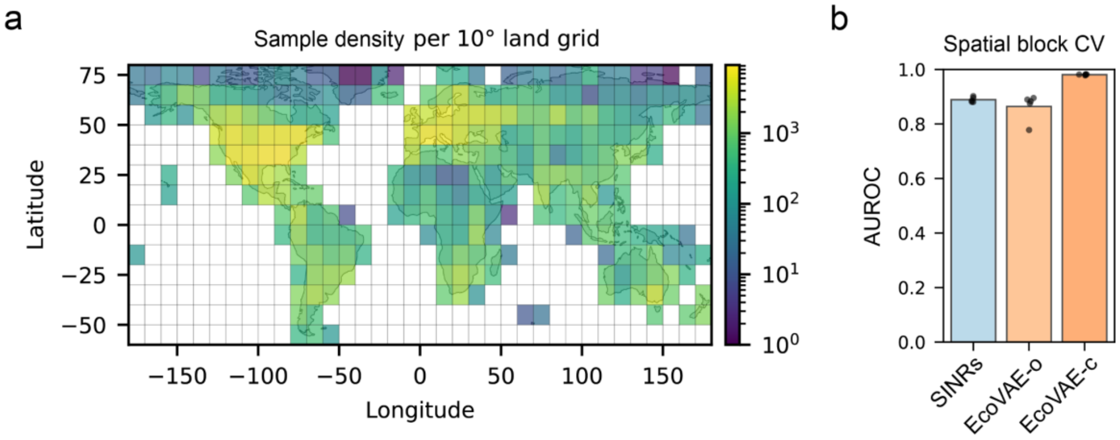

**Supplementary Figure 4. Model performance for plant distribution at the species level. a,** The distribution of AUROC for masked species in three test regions. The mean AUROCs are shown; **b,** The distribution of normalized MSE values in three test regions; **c,** Correlation between observed genera counts per grid (or observed grid counts per genus, upper panels) and predicted genera counts per grid (or predicted grid counts per genus, lower panels) across the three testing regions.

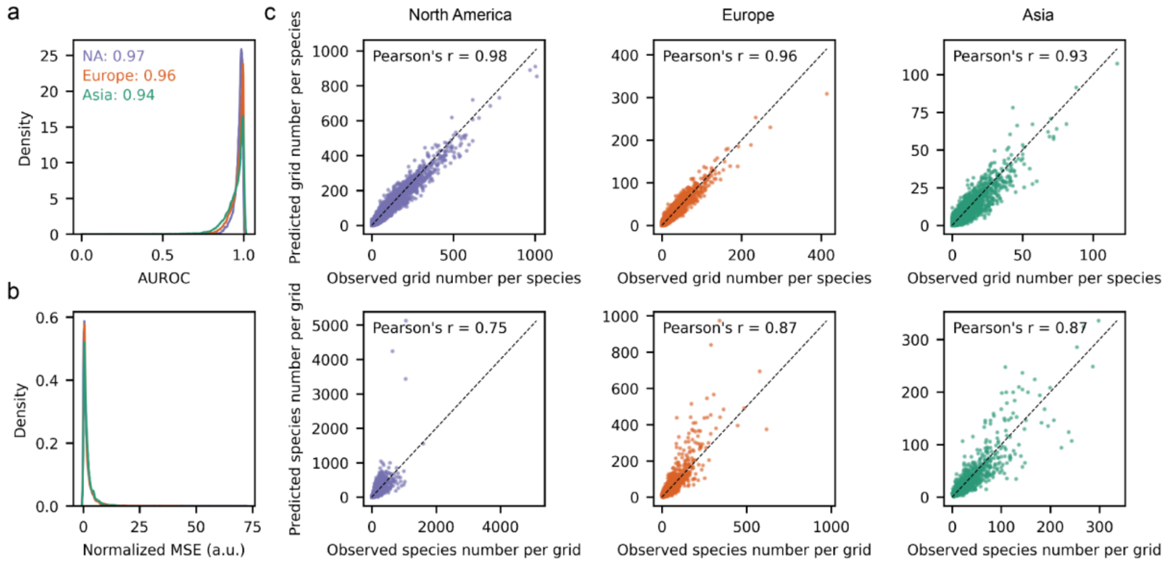

**Supplementary Figure 5. Model performance for butterfly distributions.** The distribution of AUROC for masked genera and species in three test regions are shown in **a)** and **d)**. The distribution of normalized MSE values at the genus and species levels are shown in **b)** and **e)**. **c,** Correlation between observed genera counts per grid (or observed grid counts per genus, upper panels) and predicted genera counts per grid (or predicted grid counts per genus, lower panels) across the three testing regions. **f,** Correlation between observed species counts per grid (or observed grid counts per species, upper panels) and predicted species counts per grid (or predicted grid counts per species, lower panels) across the three testing regions.

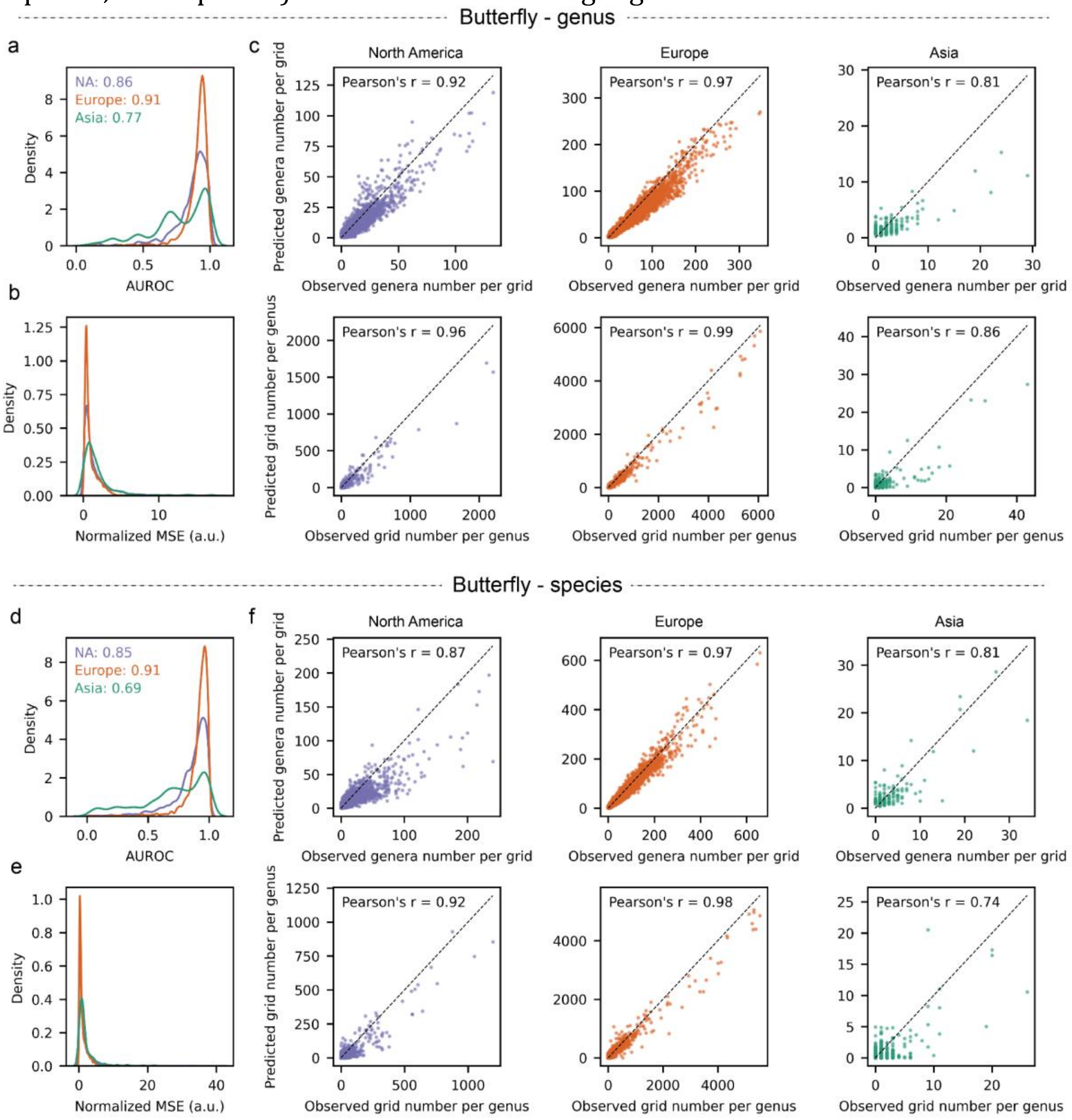

**Supplementary Figure 6. Model performance for mammal distributions.** The distribution of AUROC for masked genera and species in three test regions are shown in **a)** and **d)**. The distribution of normalized MSE values at the genus and species levels are shown in **b)** and **e)**. **c,** Correlation between observed genera counts per grid (or observed grid counts per genus, upper panels) and predicted genera counts per grid (or predicted grid counts per genus, lower panels) across the three testing regions. **f,** Correlation between observed species counts per grid (or observed grid counts per species, upper panels) and predicted species counts per grid (or predicted grid counts per species, lower panels) across the three testing regions.

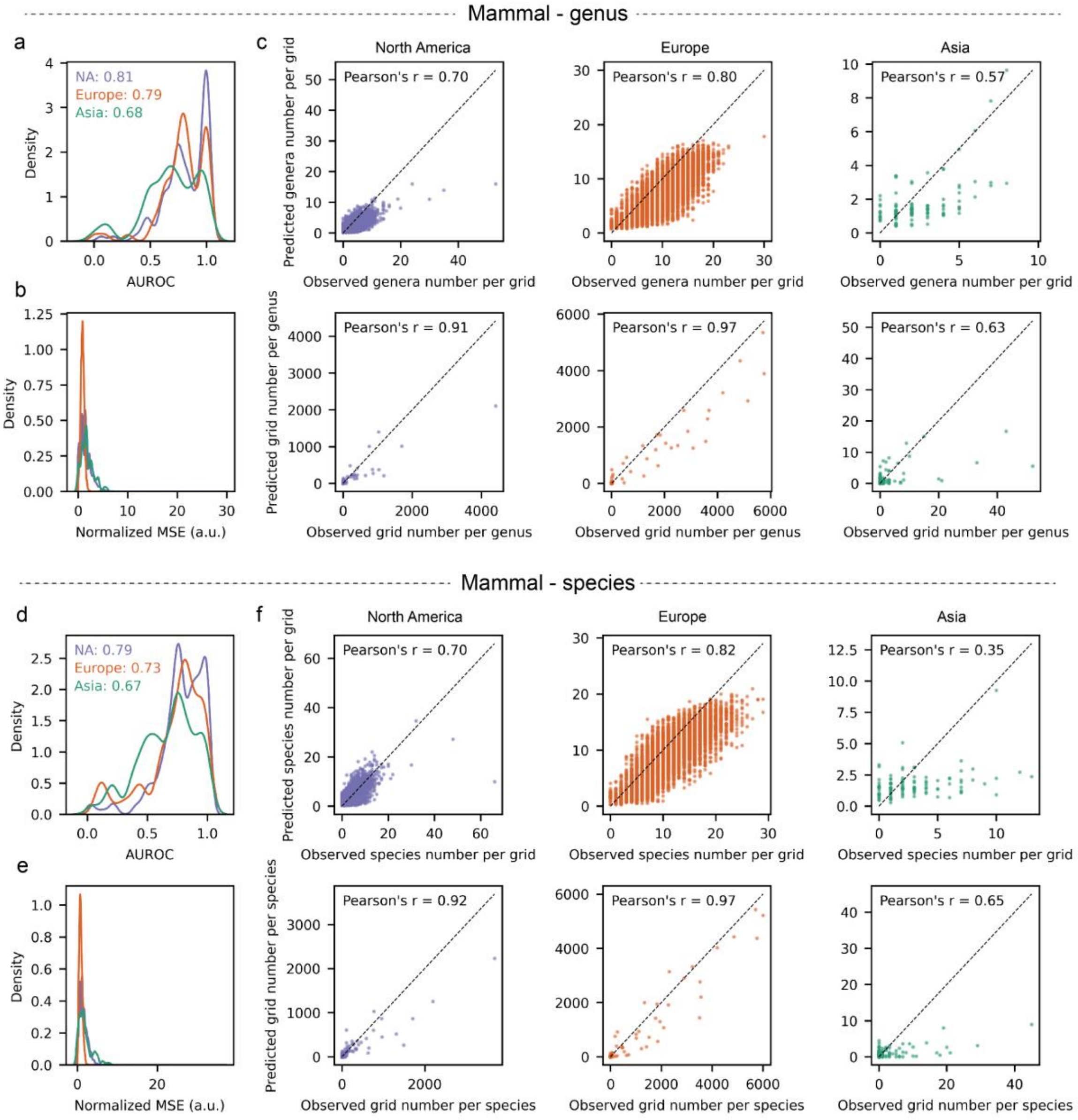

**Supplementary Figure 7**. **Comparison of GBIF observation density and prediction error. a,** Number of Global Biodiversity Information Facility (GBIF) occurrence records per 0.1° × 0.1° grid cell. **b,** Number of plant genera per 0.1° × 0.1° grid cell. **c,** Correlation between per-grid prediction error and observation count and plant genus richness across all grids, each dot represents one 0.1° × 0.1° grid cell.

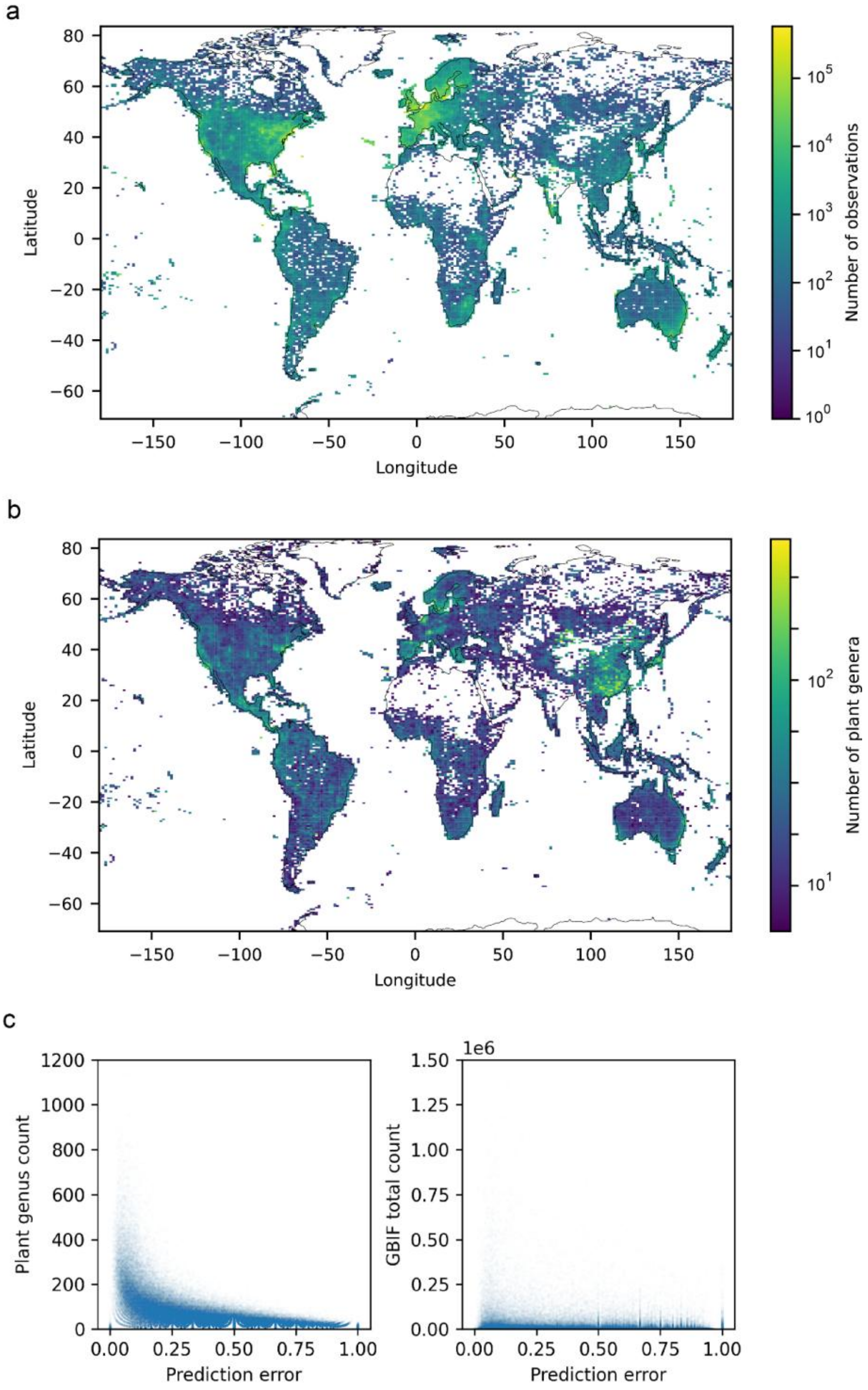

**Supplementary Figure 8. Global distribution of relative collection completeness for butterfly and mammal.** The relative completeness is represented by prediction error of EcoVAE. Darker color represents lower prediction error and higher completeness, while lighter color represents higher prediction error and lower completeness.

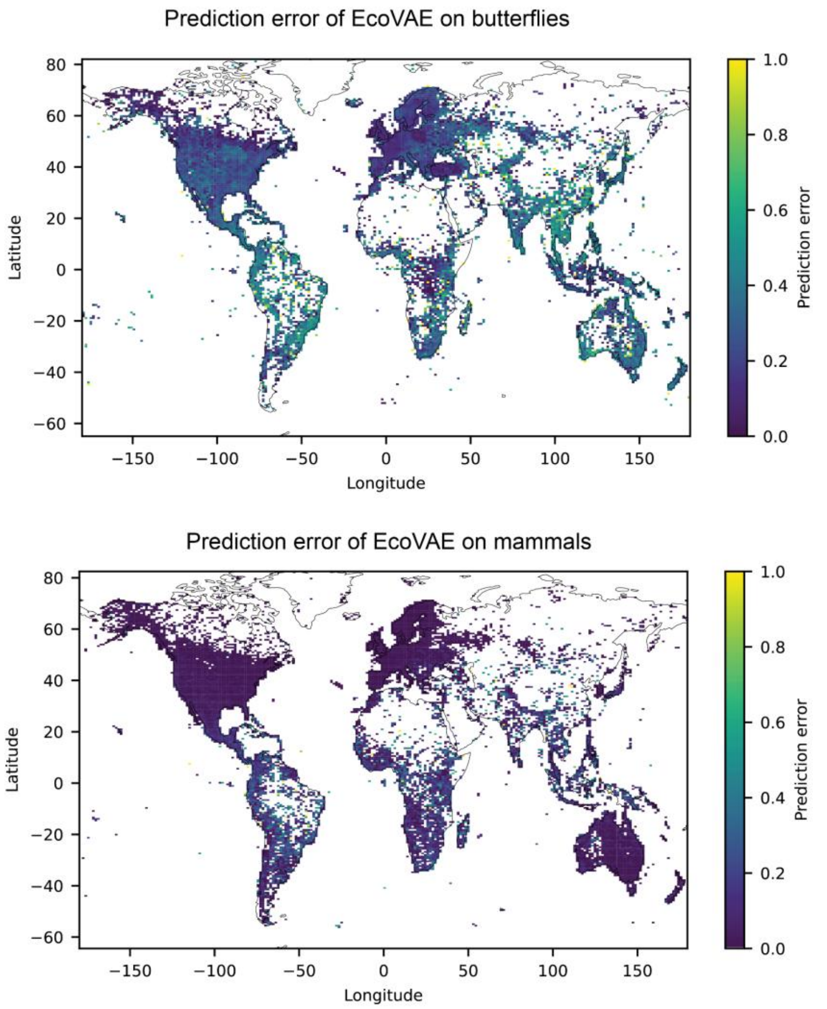

**Supplementary Figure 9. The relationship between the number of occupied grids and the overlapping index of each genus in the testing region of North America.** Each dot represents a single genus. The blue line indicates the smoothed trend, while the shaded region represents the confidence interval, both estimated using the "Loess" method. The overlapping index quantifies the degree of overlap between EvoVAE predictions and iNaturalist data, relative to a random model (see Methods for details).

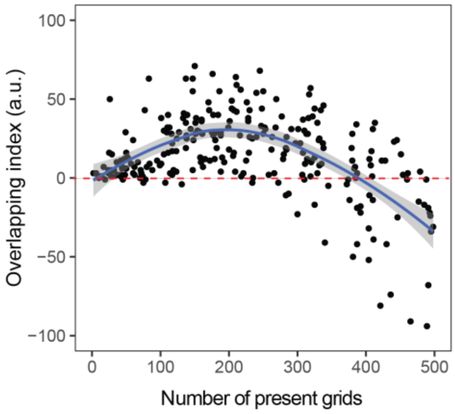

**Supplementary Figure 10. Position of the perturb testing region and distribution of biome types.** The testing region is the area inside the black rectangle. The biomes were mapped using WWF biome maps of the world (Reference: Olson, D. M., Dinerstein, E., Wikramanayake, E. D., Burgess, N. D., Powell, G. V. N., Underwood, E. C., D'Amico, J. A., Itoua, I., Strand, H. E., Morrison, J. C., Loucks, C. J., Allnutt, T. F., Ricketts, T. H., Kura, Y., Lamoreux, J. F., Wettengel, W. W., Hedao, P., Kassem, K. R. 2001. Terrestrial ecoregions of the world: a new map of life on Earth. Bioscience 51(11):933-938).

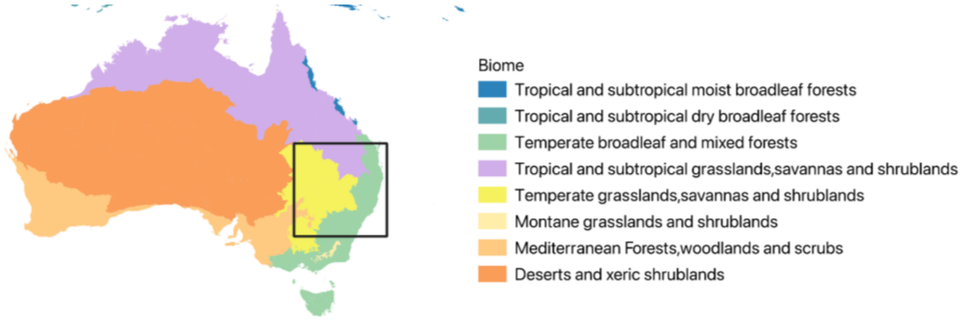

**Supplementary Figure 11. The relationship between occupied grid number of each genus and that after the addition of genus Arachniodes in the testing region.** Blue shaded area shows the 99.99% confidence interval of the linear regression. Each dot represents a single genus. Red dots represent outlier genera, i.e., the range size was significantly impacted after the addition of the focal genus.

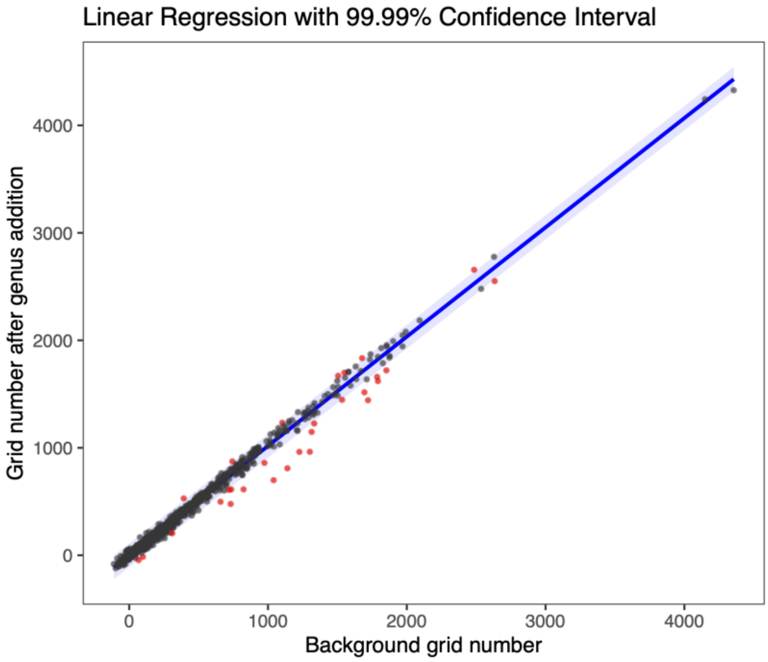

**Supplementary Figure 12. The density distribution of interactions and the relationships between genus range and interactions. a,** Density plot of the number of genera that a genus can impact; **b,** relationships between the number of grids a genus presents and the number of genera that the genus can significantly impacted; **c,** density plot of the number of genera that can significantly influence the focal genus; **d,** relationships between the number of grids a genus presents and the number of genera that can significantly influence the genus.

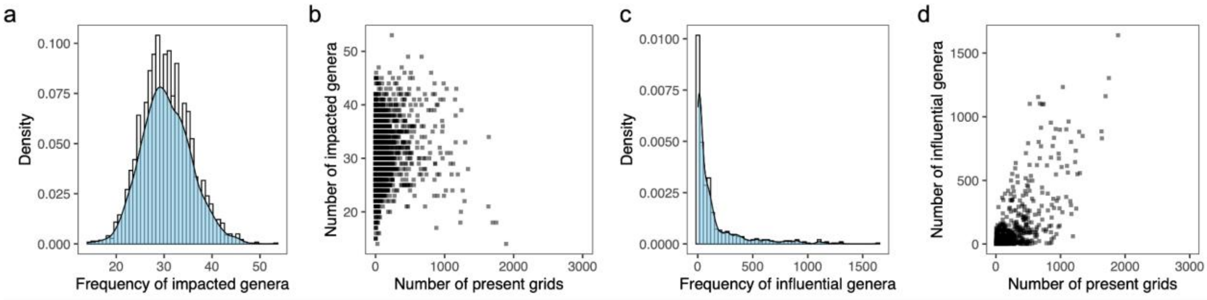

**Supplementary Figure 13. The key families with high sensitivity and influence in genus-level interactions.** The left side represents families with a high out-degree (influencing others), while the right side represents families with a high in-degree (being influenced). The width of the bands corresponds to the relative proportion of genera contributing to these interactions.

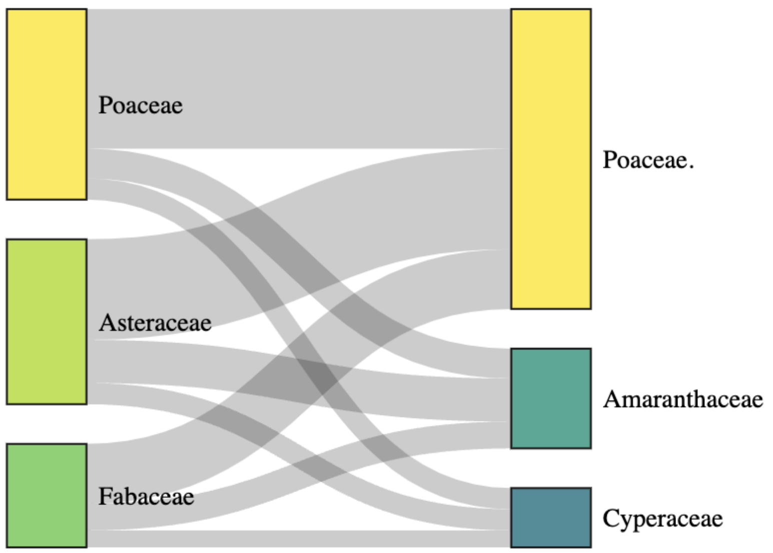
